# NOise Reduction with DIstribution Corrected (NORDIC) PCA in dMRI with complex-valued parameter-free locally low-rank processing

**DOI:** 10.1101/2020.08.25.267062

**Authors:** Steen Moeller, Pramod Kumar Pisharady, Sudhir Ramanna, Christophe Lenglet, Xiaoping Wu, Logan Dowdle, Essa Yacoub, Kamil Uğurbil, Mehmet Akçakaya

**Author notes:** Corresponding Author: Steen Moeller, Ph.D.

## Abstract

Diffusion-weighted magnetic resonance imaging (dMRI) has found great utility for a wide range of neuroscientific and clinical applications. However, high-resolution dMRI, which is required for improved delineation of fine brain structures and connectomics, is hampered by its low signal-to-noise ratio (SNR). Since dMRI relies on the acquisition of multiple different diffusion weighted images of the same anatomy, it is well-suited for denoising methods that utilize correlations across the image series to improve the apparent SNR and the subsequent data analysis. In this work, we introduce and quantitatively evaluate a comprehensive framework, NOise Reduction with Distribution Corrected (**NORDIC**) PCA method for processing dMRI. NORDIC uses low-rank modeling of g-factor-corrected complex dMRI reconstruction and non-asymptotic random matrix distributions to remove signal components which cannot be distinguished from thermal noise. The utility of the proposed framework for denoising dMRI is demonstrated on both simulations and experimental data obtained at 3 Tesla with different resolutions using human connectome project style acquisitions. The proposed framework leads to substantially enhanced quantitative performance for estimating diffusion tractography related measures and for resolving crossing fibers as compared to a conventional/state-of-the-art dMRI denoising method.

**Highlights:**

- We propose a framework, NORDIC, for denoising complex valued dMRI data using Gaussian statistics
- The effectiveness of the proposed denoising method is distinguished by the ability to remove only signal which cannot be distinguished from thermal noise
- The proposed method outperforms a state-of-art method for denoising dMRI in terms of fiber orientation dispersion
- Quantitative evaluation of NORDIC across different resolutions and SNR using human connectome type acquisitions and analysis shows up to 6 fold improvement in apparent SNR for 0.9mm whole brain dMRI at 3T.

## Introduction

Magnetic Resonance Imaging (MRI) provides a collection of different approaches that currently occupy an indispensable role in the armamentarium of methods employed for studying the human brain. Diffusion-weighted MRI (dMRI) (review (Moeller et al., 2020), and references therein), is one of these critically important techniques; it is currently the only non-invasive imaging method available to map short and long-range anatomical connections in the brain and to extract information on the white matter microstructure (Alexander et al., 2019). Complementing dMRI, there exists other MRI techniques such as resting state functional magnetic resonance imaging (rfMRI) employed for inferring functional connectivity from correlations in spontaneous temporal fluctuations (Smith et al., 2013), task based fMRI (tfMRI) that depicts regional responses to specific cognitive processes and stimuli (Barch et al., 2013), and arterial spin labeling (ASL), which is a method that provides quantitative measurements of cerebral blood flow (CBF) without the use exogenous contrast agents (Alsaedi et al., 2018). All of these methods are challenged by the inherently low signal-to-noise ratio (SNR) of the MR images themselves especially when ambitious improvements on spatial and/or temporal resolutions are sought, as foreseen, for example in the BRAIN Initiative in order to meet the enormous challenges faced in the effort to understand human brain function (Jorgenson et al., 2015).

Therefore, in all applications of MRI, particularly in the aforementioned approaches for the study of the human brain, efforts to effectively increase SNR plays a central role. Although the ultimate goal is to do so without compromising any information, attempts to do so generally trade off some other information or feature of the data, such as true spatiotemporal resolution, or specificity to the biological process of interest. This is especially evident for denoising techniques (Alkinani and El-Sakka, 2017; Fan et al., 2019; Kaur et al., 2018; Shao et al., 2014), where combinations of removing signal is balanced with corresponding feature enhancement to maintain the desired information. This is moreover challenged by most methods having to adapt to the spatially varying and non-Gaussian nature of the noise in magnitude MRI data, and the need to provide application-specific validations (Aja-Fernandez et al., 2011; Foi, 2011; Ma et al., 2020; Manjon et al., 2013).

dMRI has an inherent assumption of redundancy, since the models of the underlying biological environment has lower complexity than the amount of data acquired. The redundancy can be explicit in repetitive acquisitions, which lengthens scan time, or implicit in using a probabilistic model-fit with a lower-dimensional continuous model to a higher dimensional discrete sampling, in order to reduce the outlier sensitivity and goodness-of-fit error (Andersson and Sotiropoulos, 2015). On the other hand, denoising utilizing the well-known non-local means (A. Buades et al., 2005) was early on applied to dMRI image series (Wiest-Daessle et al., 2007), and the redundancy for local patches was first demonstrated for routine dMRI by Manjon et al. (Manjon et al., 2013) using an empirical threshold of the eigenvalues of a Principle Component Analysis (PCA) decomposition. For routine dMRI, the most advanced framework for denoising involves the Non Local Spatial and Angular Matching (NSLMA) (St-Jean et al., 2016), which incorporates Rician noise modelling, dictionary training and subspace processing. Recent work on processing complex data and adaptive to half Fourier acquisition with temporal heteroscedastic sampling, involves a low-rank spectral D-transformation using Frobenius norms, and a generalized singular value shrinkage (Cordero-Grande et al., 2019), and was shown to match the performance of NSLMA on magnitude data.

Currently, the most widespread method for suppression of noise in dMRI is the Marchenko-Pastur Principle Component Analysis (MPPCA) approach (Veraart et al., 2016), which simultaneously estimates the amount of noise and signal components in magnitude MR data adapted by using a local patch based PCA approach to essentially *remove components that have little contribution to the variance*. MPPCA uses PCA with hard thresholding on singular values, with a threshold based on asymptotic mathematical properties of random matrices (MPPCA threshold). In this approach, however, the components that have been removed is challenging to describe for a finite series with unknown low-rank.

In this work, we tackle these challenges and propose NOise Reduction with DIstribution Corrected (**NORDIC**) PCA method for reducing the influence of noise. **NORDIC** uses a dedicated processing approach to ensure that the noise component is additive with independent, identically distributed, zero-mean Gaussian entries. Using this characterization, results from random matrix theory can be efficiently used to devise a parameter-free objective threshold. For **NORDIC,** this threshold value is both numerically quantifiable and descriptive as *the removal of all components which cannot be distinguished from Gaussian noise*. **NORDIC** is similar to other PCA or low-rank based approaches, but unlike these methods, it uses *known* information from the acquisition to transform the data to fit the algorithm instead of either estimating the necessary information or adapting the algorithm to fit the data. This approach for denoising is unique from previous methods as it has negligible, if any, impact on real MR signals and can be more generally applied to different types of MRI data without re-calibration or optimization. This would in turn allow for much higher resolutions and/or reduced scan times of otherwise SNR-starved MR protocols. The cost of the method is no more than having a clean sampling of the noise and the subsequent computational requirements, both of which could be built into an online acquisition protocol and reconstruction pipeline.

In this paper, we present an extensive and quantitative evaluation of **NORDIC** on dMRI data acquired on multiple subjects with different resolutions and SNR levels. The detectability of crossing fibers in Human Connectome Project (HCP) type dMRI data (Sotiropoulos et al., 2013a) was used as the metric for assessing the performance of **NORDIC** on dMRI acquisition, since the model complexity for using this type of dMRI data is advanced and well understood. Furthermore, the increase in the ability to estimate fibers from low SNR acquisitions processed with **NORDIC** was validated by comparing it with multiple repetitions of the diffusion acquisitions that were averaged to increase the SNR.

Part of this work was presented at the International Society for Magnetic Resonance Imaging, 2017 (Moeller et al., 2017).

## Methods

### Locally Low-Rank Model

We consider the reconstructed complex-valued volumetric dMRI image series following an accelerated parallel imaging acquisition, 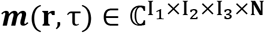 with **r** specifying the location in 3D space, and τ ∈ {1, ···, *N*}. For dMRI, *N* is the number of *q*-space samples collected with different diffusion weighting. In locally low rank (LLR) approaches, for a voxel located at **r_0_**, a *k*_1_ × *k*_2_ × *k*_3_ patch is selected whose top left corner is located at the given voxel. For ease of notation we will not explicitly write the dependence on **r_0_**, and consider arbitrary patches. For a given diffusion weighting, *τ* ∈ {1, ···, *N*} this volume is vectorized to ***y_τ_***. These vectors are then used to generate a Casorati matrix 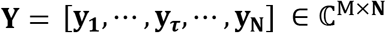, where *M* = *k*_1_*k*_2_*k*_3_. This represents the noisy data for that patch across the image series with different diffusion weighting (i.e. q space points). The denoising problem is to recover the corresponding underlying data Casorati matrix **X**, based on the following model

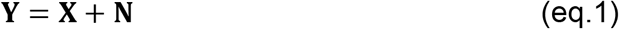

where 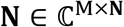 is additive Gaussian noise.

The underlying assumption for LLR models is that for any patch across the image, the data Casorati matrix **X** for that patch can be represented with a low-rank representation. Thus, LLR methods perform singular value thresholding, typically using hard or soft thresholding. Letting the singular value decomposition of **Y** be **U** · **S** · **V**^*H*^, where **U** and Vare unitary matrices, and **S** is a diagonal matrix whose diagonals are the spectrum of ordered singular values, *λ*(*j*), *j* ∈ {1, ···, *N*}. For LLR the singular values below a threshold *λ*(*j*) < *λ_thr_* are replaced by *λ*(*j*)=0 and the other singular values are either unaffected, as in hard thresholding, or reduced by *λ_thr_*, as in soft thresholding. Letting **S**_*λ_thr_*_ be the new diagonal matrix generated as a result of thresholding, the low-rank estimate of **Y** is given as ***Y_L_*** = **U** · **S**_*Λ_thr_*_ · **V**^*H*^. These locally low-rank estimates are then combined to generate the denoised image series ***m**^LLR^* (**r**, τ) either by averaging the corresponding patches together or using non-overlapping patches.

### Data-driven estimation of the threshold - MPPCA

The threshold for ***Y_L_*** can be selected empirically based on which components exhibit spatio-temporal features (Salimi-Khorshidi et al., 2014) or it can be selected based on where the noise properties mixes with the signal (Veraart et al., 2016). This latter approach is the basis for the MPPCA method, often used in dMRI and applied to magnitude dicom images, and referred to as dwidenoise (part of MRTrix, http://www.mrtrix.org) in the community. The Marchenko–Pastur law, which forms the basis for MPPCA, describes the asymptotic properties of singular values in random matrices with independent identically distributed (i.i.d.) zero-mean entries. For such a random matrix **Z** of dimension *M* × *N* with *M* > *N* and let 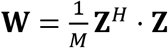. The spectrum of **W** is band-limited as a function of the variance of the entries and the matrix dimensions in the asymptotic limit. Specifically, for a matrix **Z** with i.i.d. entries having mean 0 and variance *σ*^2^, the singular values of **W** are asymptotically band limited to values between *λ*_-_ and *λ*_+_, where 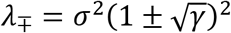 with 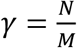, i.e. the spectrum has a bandwidth 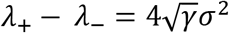.

The MPPCA leverages this distribution to select the threshold for denoising, where the Casorati matrix has both signal and random noise contributions. In this case, the tail of the spectrum of **S** is used to simultaneously estimate the noise level, and the value of *λ_thr_*. For a fixed value of signal components, the width of the tail, assuming zero-mean i.i.d. noise, satisfies with high probability the asymptotic limit

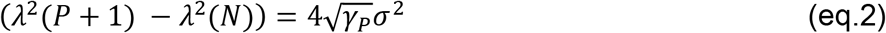

where *γ_P_* = (*N – P*)/*M* while also satisfying the inequality

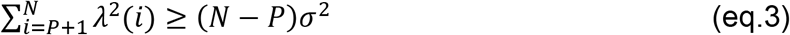

The band limiting of the spectrum, allows for calculating the largest value of *P* for which the equality (eq.3) holds, while simultaneously providing a value for the variance *σ*^2^. The use of the Marchenko-Pastur assumes i.i.d. zero-mean noise, which is typically not satisfied in reconstructed MRI data, where the noise is spatially varying from the use of undersampled k-space acquisitions. Furthermore, the equality assumes zero-mean entries, whereas the noise in magnitude images in MRI is either Rician or non-central Chi^2 distributed.

### Proposed LLR Denoising

In **NORDIC**, the data matrix **Y** is constructed so that the noise matrix component matches the random matrix theory model. This is achieved by: 1) retaining the images as complex valued with zero-mean Gaussian noise following image reconstruction, 2) mapping the spatially varying noise in the reconstructed images to spatially identical noise using the g-factor of the parallel imaging method, and 3) selecting the threshold explicitly based on the noise spectrum.

For the first step, we utilize slice-GRAPPA reconstruction for the slice accelerated dataset, obtained with the simultaneous multislice (SMS)/Multiband (MB) approach (Moeller et al., 2020). A single kernel 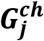 is constructed for SMS/MB with phase-encoding undersampling such that for each slice, *j*, and channel, *ch*,

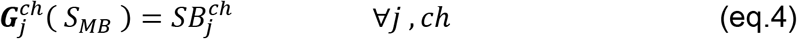

and the kernels 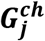 are calculated similarly as in slice-GRAPPA from the measured individual slices *SB_i_* with 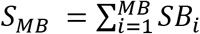. For the combination of reconstructed individual channels, the SENSE-1 reconstruction (Sotiropoulos et al., 2013b) is used to maintain Gaussian noise (Aja-Fernández and Vegas-Sánchez-Ferrero, 2016) in the complex valued reconstructions and the ESPIRIT algorithm (implemented from the Berkeley Advanced Reconstruction Toolbox (BART) https://mrirecon.github.io/bart/) is used for sensitivity estimation.

For the second step, g-factors are calculated building on the approach outlined in (Breuer et al., 2009) for g-factor quantification in GRAPPA reconstructions and detailed in (Moeller et al., 2020) and the same ESPIRIT sensitivity profiles used for image reconstructions are also used for the determination of the quantitative g-factor. The g-factor is subsequently used to normalize the signal scaling in ***m***(**r**, τ), as ***m***(**r**, τ)/*g*(**r**). We note that this ensures zero-mean and spatially identical noise in a given patch. The remaining independence requirement is satisfied by choosing the patch size small enough to ensure that no two voxels within the patch are unaliased from the same acquired data, which can be done by selecting *k*_3_ < *I*_3_/*MB* and *k*_2_ < I_2_/*R*, where *MB* is the acceleration rate (i.e. the number of simultaneously excited slices) along the slice direction using RF pulses with MB number of bands, and, *R*, the in-plane phase-encoding undersampling rate. When blipped-CAIPI (Setsompop et al., 2012) encoding is added, then the patches in the MB unaliased slices do not overlap directly and *k*_3_ > I_3_/*MB* can be used, but is dependent on the FOV shift and the *R* factor.

Following these steps, the noise component of **Y** has zero-mean i.i.d. entries, and the threshold in the ideal setting is given as the first singular value specified by the Marchenko-Pastur law for such *M* × *N* noise matrices. This choice ensures that all components that are indistinguishable from Gaussian noise are removed. While this threshold can be calculated from the analytical formula, this is an asymptotic expression, and deviations may occur for the practical finite matrix case. Thus, for a finite-sized random matrix, an alternative is needed in the absence of an analytical expression and we calculated this threshold by using noise images (where no RF excitation is applied), which were still reconstructed identically to the acquired data, and also corrected with the g-factor. We use a Monte-Carlo simulation with matrices of size *M* × *N* extracted from the noise image to generate the sample average for the largest singular value of an *M* × *N* random matrix with i.i.d. zero-mean elements and variance *σ*^2^ identical to the noise images, and use this as the threshold *λ_thr_*.

Since the g-factor normalization changes the signal scaling, after **NORDIC** processing, the volumes with noise reduction applied ***m^N0RDIC^***(**r**,τ) are further processed as ***m^NORDiC^***(**r**, τ) · *g*(**r**) such that the signal magnitude is corrected back to the original form. A schematic of the steps in the proposed **NORDIC** algorithm is shown in **Figure 1**.

**Figure 1.**
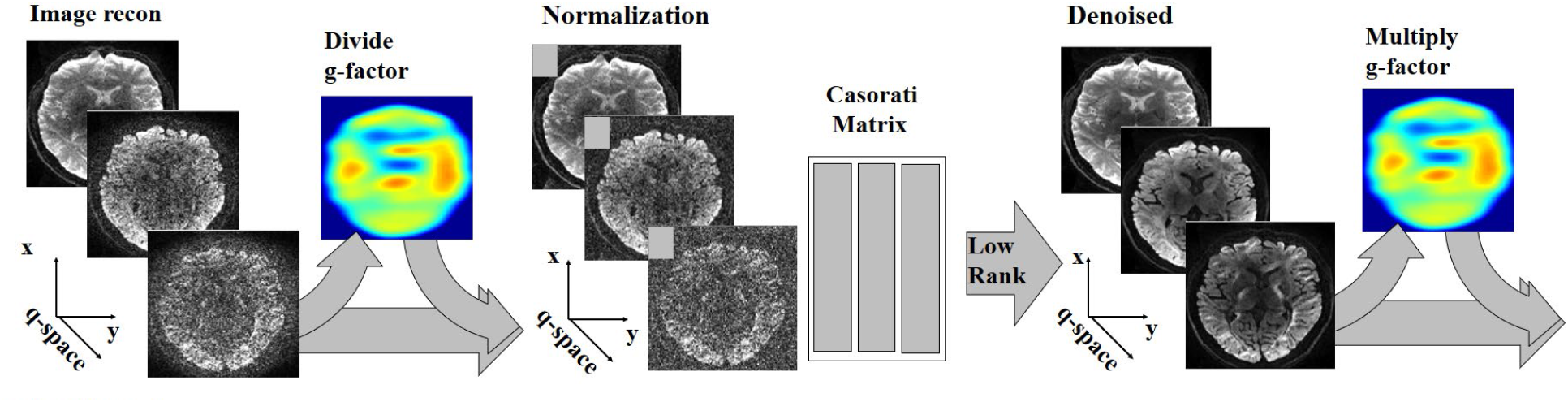
Flowchart of the **NORDIC** algorithm for a series ***m***(**r**, τ). Firstly the series is normalized with the calculated g-factor kernels as ***m***(**r**, τ)/*g*(**r**). From the normalized series the Cassorati matrix **Y** = [**y**_1_, ···, ***y_j_***, ···, **y_N_**] is established and the low-rank representation of ***Y*** is calculated as ***Y_L_*** = **U** · **S***_λ_thr__* · **V**^*H*^, where *λ*(*i*) = 0 for *λ*(*i*) < *λ_thr_*. After reforming the series ***m^LLR^***(**r**, τ) the normalization with the calculated g-factor is reversed as ***m^NORDIC^***(**r**, τ) = ***m^LLR^***(**r**, τ) · *g*(**r**).

### Patch averaging

The transformation from patches to a Casorati matrix removes an explicit spatial connection between rows, which is re-established when the processed Casorati matrix is reordered to a patch. The selection of patches can be anywhere between non-overlapping and maximally overlapping, directly proportional to an increase in computation time. With non-overlapping patches remaining block-artifacts are commonly observed. For patch averaging, the patches are used independently of the *x,y,z* orientations, i.e. independent of how they are acquired in terms of readout, phase-encoding and slice-encoding direction (Katkovnik Vladimir et al., 2010). For the over-lapping patches, the combination of these can be weighted equally, as used here, or weighted as the number of retained components, which was introduced for earlier PCA methods (Ma et al., 2020) with negligible differences when a uniform threshold is used.

### (*x* + *t*) phase-stabilization

The use of complex valued information is in itself nothing ominous for LLR techniques. For dMRI, the phase induced in the process of diffusion weighting, the diffusion phase, is caused by the temporal fluctuations of the B0 field over the head during diffusion encoding primarily due to respiration (Anderson and Gore, 1994), and is immaterial in the dMRI modelling, which only considers the changes in signal magnitude subsequent to diffusion encoding for accessing underlying tissue properties. However, keeping the diffusion phase increases the number of components necessary to describe the signal. The appearance of the diffusion phase can be reduced, based on the fact that the phase is independent of tissue properties and is spatially smooth, while also noting that it can have 2π phase-wraps. For a dMRI series with different q-samples, the volume and slice specific smooth phase is removed in a two-step process where first the common phase per slice is removed (using an average over all volumes), and then the volume specific smooth phase is removed as follows. Each volume is Fourier transformed and multiplied with a 2D weighted Tukey filter (the outer product of two weighted Tukey filters of length N1 and N2 respectively, where *N*_1_, *N*_2_ are the image dimensions) which is equivalent to a broader blurring function in image space, and then subsequently transformed into image space with an inverse Fourier transform. The resulting phase, per slice and volume, is used as a low-pass filtered estimate and multiplied with the original data with the common phase removed. The effect of the 2-step phase-correction is illustrated in Supplemental Figure S1.

### Numerical Evaluation of Threshold Choice

Existing low-rank denoising methods, such as MPPCA, and the **NORDIC** approach are denoising methods that work directly on the spectrum of the Casorati matrix. Even without the considerations of the i.i.d. zero-mean entries for the noise component of the Casorati matrix, there are differences between the existing, previously published (Veraart et al., 2016), data-driven choices for the threshold and our proposed fixed threshold. To highlight these differences, a numerical simulation was performed, where the noisy data was generated according to (eq.1). The underlying noise matrix **N** was generated as i.i.d. Gaussian noise, whereas the underlying data matrix **X** was generated using two random unitary matrices **U**_X_ and **V**_X_, and a low-rank spectrum **S**_X_. We use *λ*_**N**_ to denote the spectrum of the noise matrix **N**, and 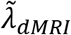 to denote the spectrum extracted from a 1.5mm isotropic resolution Lifespan dMRI acquisition (Harms et al., 2018) with 99 *q*-space samples, described further below. These are scaled individually relative to the spectra *λ*_**N**_ of the noise, and computed for 99 volumes with 7^3^ and 11^3^ voxels respectively. Four different spectra *λ_Model_, λ_LR Model_, λ_dMRI_* and *λ_LR dMRI_* were considered for **S**_X_ with

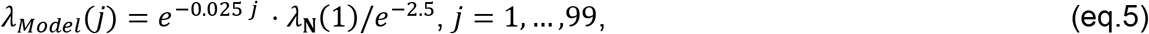

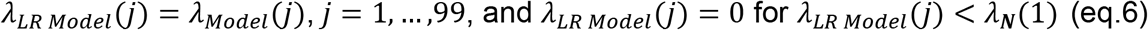

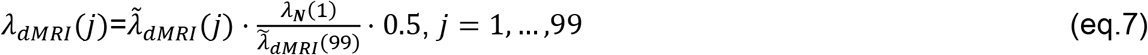

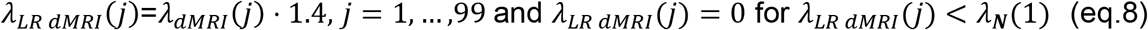

The constants in (eq. 5) – (eq. 8) do not correspond to a specific acquisition bio-physical model, and *λ_Model_, λ_LR Model_, λ_dMRI_* and *λ_LR dMRI_* are considered in terms of their effect for a numerical simulation of the techniques.

Subsequently, the singular value spectra of N, X and Y were calculated, and the two different threshold selection strategies were compared.

### In-vivo imaging

Data were acquired at the Center for Magnetic Resonance Research (CMRR), University of Minnesota (UMN). All participants provided written informed consent and the study was approved by the UMN’s Institutional Review Board. Eight participants were scanned on a Siemens Magnetom Prisma (Siemens Healthcare, Erlangen, Germany) 3 Tesla (3T) scanner equipped with a 32 channel head coil and a 80 mT/m gradient system with a slew rate of 200 T/m/s. An additional participant was scanned on a 7 Tesla (7T) MR scanner (Siemens, Erlangen, Germany) equipped with 32 receive channels and the Siemens SC72 body gradient that achieves 70 mT/m maximum strength and 200 T/m/s maximum slew rate with the current gradient drivers; maximum slew rate usable for diffusion encoding gradients was, however, limited to ~125 T/m/s due to peripheral nerve stimulation (Vu et al., 2015). The vendor supplied Nova single-channel transmit and 32-channel receive head coil was employed for RF transmission and signal reception. The data acquisition and image reconstruction was performed with the CMRR distributed C2P multiband diffusion sequence (https://www.cmrr.umn.edu/multiband/) on both systems.

Data were acquired at 3T where three different nominal isotropic imaging resolutions of 1.5mm, 1.17mm and 0.9mm were used, leading to effective voxel volume ratios of 1:0.5:0.2. A two-shell diffusion sampling scheme was employed with *b*=1500, 3000 s/mm^2^ with 99 *q*-space directions (46 for b=1500 s/mm^2^, 46 for b=3000 s/mm^2^ and 7 for b=0 s/mm^2^, as an interleaved combined set following Caruyer (Caruyer et al., 2013)); these data were acquired twice, running the phase encode direction either in the AP or PA directions, as in the HCP (Sotiropoulos et al., 2013a; Ugurbil et al., 2013). We followed the HCP Lifespan protocol (Harms et al., 2018) for each AP or PA acquisition; this 1.5mm resolution dMRI protocol deviated from the HCP Lifespan protocol in one aspect, namely the HCP acquired such data twice (Harms et al., 2018) whereas in our case this was done only once, collecting as a result half the data compared to the HCP Lifespan protocol. For the higher resolution acquisitions, by necessity, we deviated somewhat more from the HCP Lifespan protocol; in this case, for each resolution, the *q*-space sampling was the same as that employed for the 1.5mm acquisition but the acceleration factors changed in order to keep the TR approximately in the same range. In all cases, the slice-orientation was chosen similarly to the HCP as oblique coronal-axial along the AC-PC line to reduce the necessary number of slices to cover the whole brain. Each resolution was acquired with 6/8 partial Fourier (similar to the Lifespan and the HCP-young adult (Sotiropoulos et al., 2013a; Ugurbil et al., 2013)) and was acquired with the following parameters:

<u>1.5mm isotropic</u>: TE/TR=89.2/3230ms, MB×R=4×1, echo-spacing 690μs; 92 slices
<u>1.17mm isotropic</u>: TE/TR=77.8/2780ms, MB×R =5×2, echo-spacing 770μs; 120 slices
<u>0.9mm isotropic</u>: TE/TR=95.4/5850ms, MB×R =4×2, echo-spacing 940μs; 152 slices.

The highest and lowest spatial resolutions were chosen in accordance with the restrictions imposed by gradient resonance frequencies. For 5 subjects, all three resolutions were obtained. For the remaining 3 subjects 60min of data were collected at one of the three resolutions. The 60-min data acquisition time allowed 6, 5, and 3 repeated acquisitions, respectively for the 1.5mm, 1.17mm and the 0.9mm data.

As single whole brain dMRI data set acquired at a nominal 0.7mm isotropic resolution was also evaluated for denoising. This data set was obtained at 7T, with MB×R=2×3, TR/TE=13.4s/91ms and partial Fourier of 6/8. The FOV was 208×208×126mm and 180 oblique axial slices was used to cover the whole brain. A two-shell diffusion sampling scheme was employed with *b*=1000, 2000 s/mm^2^ having total 96 interleaved *q*-space directions and an additional 11 for b=0 s/mm^2^. These data were acquired twice, running the phase encode direction either in the AP or PA directions, and obtained as 4 independent acquisitions of ~15min duration, including the auto-calibration and single band reference scans. All data were processed as magnitude data and corrected for motion, geometric and eddy currents induced distortions and outliers with FSL TOPUP (Andersson et al., 2003) and EDDY (Andersson et al., 2016; Andersson and Sotiropoulos, 2016). For repeated acquisitions, each individual series was processed independently, and all were subsequently motion corrected and averaged. DTI and multi-shell crossing fibers models were fitted to the corrected data, using DTIFIT and BEDPOSTX with a multi-exponential decay assumption (Jbabdi et al., 2012), respectively, and results were visualized with FSLEyes.

### In-Vivo Imaging Evaluation of NORDIC

#### Qualitative Evaluation

A real-valued simulation was performed to evaluate the proposed **NORDIC** method using a high SNR reference volume with spatial matrix size 140×140×92, and with 99 volumes (*q*-space samples). The reference was the EDDY corrected (Andersson et al., 2016; Andersson and Sotiropoulos, 2016) standard reconstruction data from 6 averages for the 1.5 mm resolution listed in the “In-vivo imaging” section. This high SNR reference was degraded by adding real-valued noise. The images were evaluated in two different ways; 1) using the simulated “ground truth” real-valued images and 2) using the magnitude images to include the Rician noise properties. The noise level was selected such that for the volumes with the lowest signal intensity, the mean signal over the whole brain was 1 (with a 10:1 maximum to mean ratio), and the noise was independently and identically distributed (i.i.d.) Gaussian with variance 1. Subsequently, the proposed **NORDIC** approach was compared to MPPCA denoising.

For the afore described simulation using a high SNR dMRI data degraded by addition of noise, the impact on apparent image SNR was evaluated for **NORDIC** using different patch-sizes *M*, and compared with MPPCA using both Gaussian and Rician noise. In addition a diffusion contrast image defined as the difference between two images with different *q*-vectors directions and the same b-value,

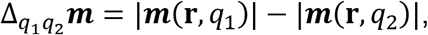

was used to evaluate blurring between different q-space acquisitions from the low-rank processing. The two *q*-vectors have the same b-value, and are selected as sequential in the *q*-space set defined following Caruyer (Caruyer et al., 2013), and as such nearly orthogonal.

In order to demonstrate the whole brain effect of improved detection of crossing fibers in the **NORDIC** processed data, the connectivity of the posterior corona radiata (PCR, both left and right) with the rest of the brain was investigated qualitatively using probabilistic tractography (‘probtrackx’)(Behrens et al., 2007).

#### Quantitative evaluation

Detection of second and third fiber orientations, necessary for resolving white matter crossing fibers, is widely used as a measure of information content in the data. As such, two VOIs, each covering the superior longitudinal fasciculus (SLF) and posterior corona radiata (PCR), which contain various crossing fibers configurations, were selected for quantifying the effect on information content in the denoised data. These VOIs were defined using the JHU-ICBM 1mm atlas (https://identifiers.org/neurovault.image:1401) and were linearly transformed into the 1.5mm data space first, and then from there into the 1.17mm and 0.9mm data spaces using linear transformations for each subject. The SLF and PCR VOIs from the left and right hemispheres were combined into a single VOI for the SLF and PCR respectively. In BedpostX, automatic relevance determination (ARD) was used to accurately recover fiber orientations supported by the data during the data-driven parameter estimation process. The parameters (e.g. fiber volume fractions) that are not supported by the data will have a value around zero with very low variance in the posterior distribution.

##### fiber detection rate

The number of voxels with two- and three-way fiber crossings (referred to as second and third fibers) were normalized by the total number of atlas-based voxels in the VOI for each dataset, and reported as a percentage as a proxy for sensitivity. The normalization by the total number of voxels in the VOI compensates for the variations in the size of the VOI between different datasets and resolutions.

##### fiber orientation dispersion

The fiber orientation dispersion is a measure of the consistency in the ball and stick model based predictions of the fiber orientations and a proxy for consistency and indirectly specificity. Through estimates from Markov Chain Monte Carlo (MCMC) sampling of the posterior distribution of fiber orientation in the Bayesian estimation process in bedpostx an uncertainty can be calculated. The uncertainty in the fiber orientation estimation represents the variance in the MCMC sample orientation vectors around the distribution mean and is the *fiber orientation dispersion*. The reported *fiber orientation dispersion* is calculated as (1-s), where s is the largest eigenvalue of the average tensor constructed from these MCMC samples (Behrens et al., 2007). This calculation is done separately for each fiber populations. Higher information content in the data can result in more consistent predictions of the orientations, representing higher accuracy of estimated fiber orientation.

##### gain in fiber orientation accuracy

The gain in accuracy is the improvement (i.e. a *decrease*) in uncertainty calculated as the ratio of the dispersion of the reference acquisition ***Y*** divided by the dispersion for the target ***Y_L_***, as a proxy for gain in SNR.

Several experiments were performed to evaluate the various steps in the proposed **NORDIC** algorithm. Apart from the simulations described earlier, the necessary and additional steps of (*x* + *t*) phase-stabilization and patch averaging were evaluated for the lowest SNR data for the in-vivo acquisition.

Subsequently, the efficacy of **NORDIC** was qualitatively and quantitatively evaluated using diffusion metrics for multiple subjects at 3 different resolutions (and thus SNR levels), including FA maps, crossing fiber detection rate, fiber orientation dispersion and gain in accuracy. The impact on apparent SNR was evaluated by comparing the **NORDIC-**processed dMRI acquisitions, with high SNR data obtained by averaging repeated acquisitions. The performance of MPPCA and **NORDIC** were quantitatively compared, for multiple subjects and resolutions, in the superior longitudinal fasciculus (SLF) and posterior corona radiata (PCR) which are regions with known crossing fibers. Finally, implications for whole brain tractography across the different resolutions were qualitatively investigated using the connection strength, defined as the number of streamlines passing through each voxel connecting the seed VOIs to the rest of the brain (Behrens et al., 2007).

## Results

### Numerical Evaluation of Threshold Choice

For an ideal observation model **X**, the spectra for *λ_Model_, λ_dMRI_* and *λ_LR dMRI_* are illustrated in each column in Figure 2, respectively, along with the spectra for the observation **Y**. The top row shows the spectra for *M*=7^3^ and the bottom row show it for *M*=11^3^, both with *N*=99. For each model, the corresponding spectra for **N, X** and **Y**, along with the threshold estimated from the observation **Y** based on the MPPCA method, as well as the maximal singular value of the noise which is the threshold choice in **NORDIC**, are shown with dotted horizontal yellow and green lines respectively.

**Figure 2.**
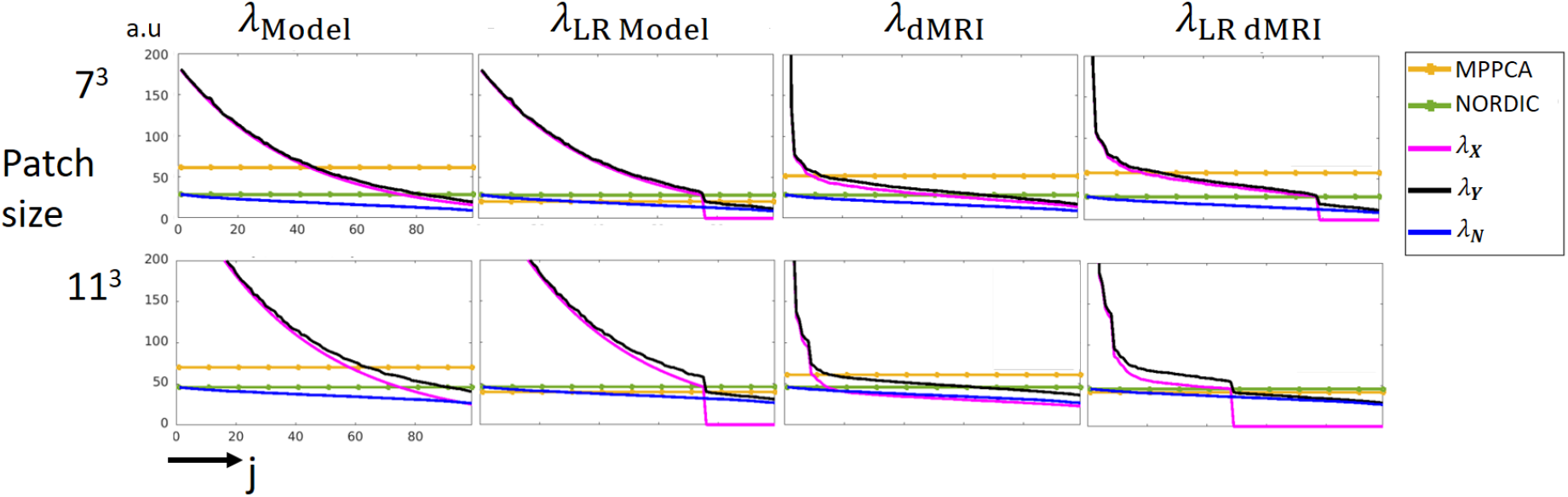
The interaction between the spectra of the underlying model, additive noise and the observed noise perturbed measurements, and the threshold estimated with asymptotic properties and hard thresholding based on the maximal singular value. The four spectra shown in each of the four columns are generated by eq.5, eq.6, eq.7, and eq.8, respectively, given in section titled Numerical Evaluation of Threshold Choice; they represent an asymptotic model (*λ_Model_*), an asymptotic model with low-rank (*λ_LR Model_*), a spectra from dMRI which is full rank and falls below the maximal singular value of the noise (*λ_dMRI_*) and a low-rank signal which does not fall below the maximal singular value of the noise (*Λ_LR dMRI_*). The MPPCA technique uses the asymptotic properties of the noise spectra to infer a threshold, and the NORDIC uses the prior knowledge of the noise-level for this.

For *λ_Model_* shown in the first column in **Figure 2**, the top row shows that the spectrum for the observation ***Y*** is close to *λ_Model_*, and for the larger patch shown in the bottom row a difference in the spectra of ***Y*** and *λ_Model_* can be noted. For both patch-sizes the MPPCA estimated transition between signal and noise components, removes more components that what is retained with **NORDIC**. For *λ_LR Model_* shown in the second column in **Figure 2**, the top row shows that the spectrum for the observation ***Y*** is close to *λ_LR Model_* for all values larger than the largest value in the spectrum of the noise indicated with the horizontal green line. For the larger patch shown in the bottom row there is a larger difference between the spectra of ***Y*** and *λ_LR Model_*. For the values below the green line, the spectrum for ***Y*** for both patch sizes has a steeper slope compared with *λ_**N**_* for the noise, and the MPPCA identifies the discontinuity in the spectrum. In this idealized low-rank scenario MPPCA and **NORDIC** behave similarly.

The third column shows results for *λ_dMRI_* which is not inherently low-rank. In the top row the spectrum for the observation **Y** is close to *λ_dMRI_*, and the MPPCA determines a threshold for *λ_dMRI_* which has fever singular values as compared with the threshold for **NORDIC**. Both thresholds reduces the number of components relative to the true *λ_dMRI_*, while MPPCA removes more components than when the threshold is applied to *λ_dMRI_*, for **NORDIC** more components are kept in the spectrum of ***Y*** than when the threshold is applied directly to *λ_dMRI_*. In the right column, for *λ_LR dMRI_*, the threshold for MPPCA in the top row finds fewer significant components as compared with the components in *λ_LR dMRI_*, whereas **NORDIC** determines the correct number. For the larger patch shown in the bottom row the number of retained components with both MPPCA and **NORDIC** match to the content in *λ_LR dMRI_*.

Figure 2 shows a numerical example of low rank thresholding for a fixed noise-level. For models with an underlying low-rank spectrum, as in column 2, Figure 2, both MPPCA and NORDIC may estimate a similar threshold whereas for the *λ_LR dMRI_* spectrum (right most column, Figure 2), MPPCA estimates a similar threshold to the spectrum for *λ_dMRI_* (third column from left, Figure 2). An increase in the patch-size can increase the spread of the singular-values between *λ_X_* and *λ_Y_*. For a model with Gaussian noise, and for spectra with the chosen decay-rates, the MPPCA may perform similar to NORDIC with a 11^3^ patch for signal that are low-rank, whereas NORDIC may retain more signal components than MPPCA for the spectrum that are not low-rank. We note these observations are for numerical purposes only, and their applicability do not necessarily extend to the *in vivo* dMRI data, for which the patch size was optimized to 5^3^ for MPPCA in (Veraart et al., 2016). Accordingly, the MPPCA processed images presented in this paper were all processed with a patch size of 5^3^.

### In-vivo Simulation

#### Qualitative comparison

**Figure 3** illustrates evaluations from simulations performed using dMRI data with 99 *q*-space samples obtained in the human brain and subsequently degraded by adding noise to it. Results from this simulations are shown for a single slice, processing it with **NORDIC** using patch sizes of 5^3^, 7^3^, and 11^3^ (Fig.3A.iii, 3A.iv, and 3A.v, respectively) and with MPPCA (Fig.3A.vi, and 3A.vii); the latter was applied both on the Gaussian noise data, and on data with Rician noise obtained by using the magnitude of the in-vivo data with added simulated noise. The slice used for these illustrations generated with the standard reconstruction is shown before and after the addition of noise in Fig.3A.i and Fig.3A.ii, respectively. Similar analysis are displayed in **Figure 3B** for the *q*-space contrast, Δ_*q_1_*q*_2_*_***m***, the difference between two images with two different *q*-vectors and same b-value. In supplemental Figure S2, the residual between the denoised and the noise degraded reference is shown.

**Figure 3.**
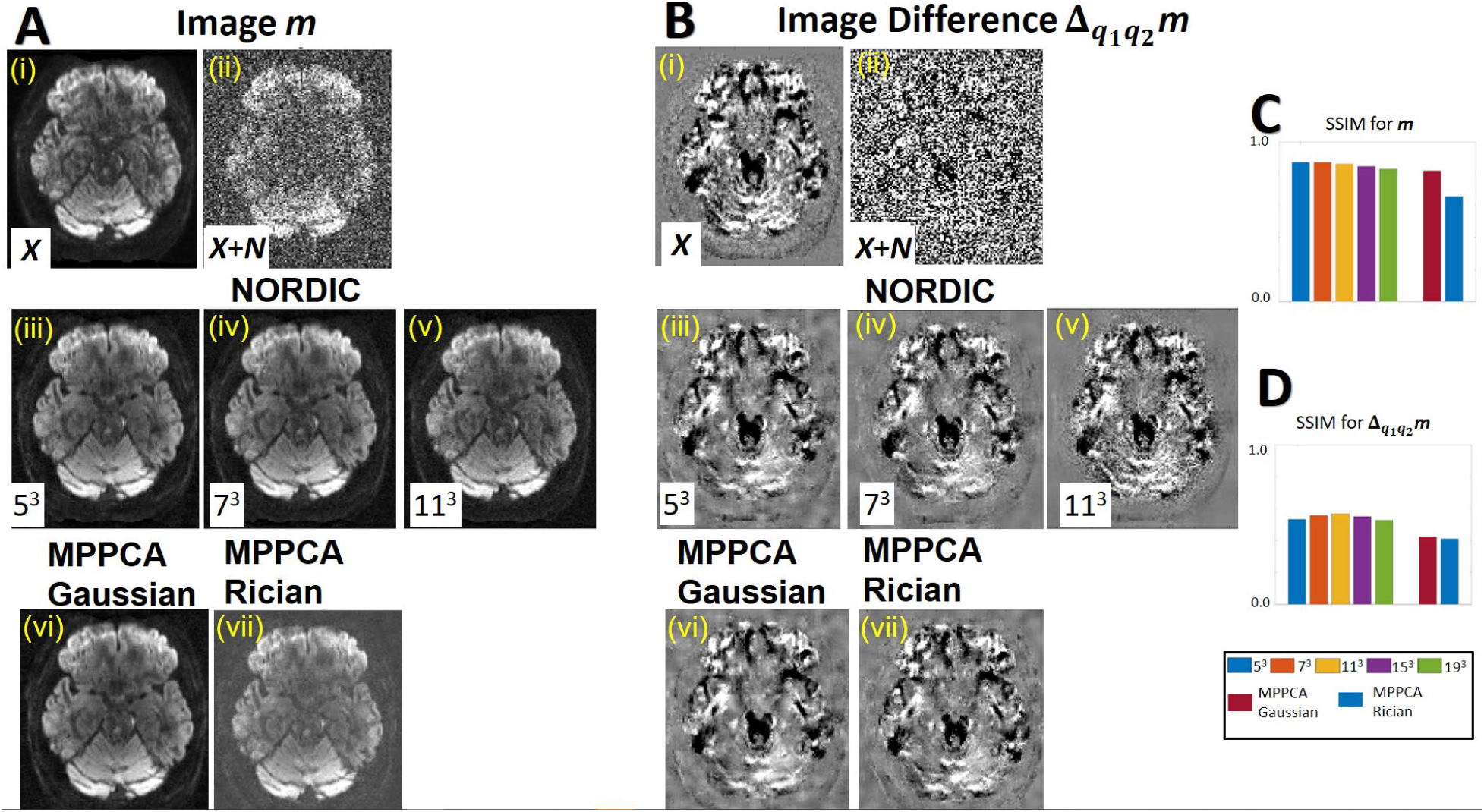
Real valued simulation of the quality of the “optimal” signal recovery with **NORDIC. Figure A** shows for a single slice the quality of the images reconstructed with denoising using NORDIC and MPPCA subsequent to SNR degradation with the addition of noise, and **Figure B** shows analogous images for *q*-space contrast, Δ_*q*_1_*q*_2__***m***, as the difference between volumes with different *q*-vectors and same b-value. The reference images are shown in Fig. 3A.i, and 3B.i before addition of noise and in Fig.3A.ii, and 3B.ii after addition of noise. The NORDIC methods are compared for patch-sizes of 5^3^, 7^3^, and 11^3^ (middle row), and the MPPCA method (bottom row) are compared using the Gaussian noise and Rician noise. Panel C shows the structural similarity index (SSIM) restricted to the brain between *m* without noise, and NORDIC with patch sizes 5^3^, 7^3^, 11^3^, 15^3^, and 19^3^. **Panel D** shows the structural similarity index (SSIM) restricted to the brain between Δ_*q*_1_*q*_2__***m*** without added noise, and NORDIC processing with patch sizes 5^3^, 7^3^, 11^3^, 15^3^, and 19^3^. The SSIM for Panels C and D are averaged over all slices and diffusion direction for the different patch sizes. The SSIM from D is 0.54 for NORDIC using a patch size of 11^3^, 0.49 for MPPCA with Gaussian noise, and 0.50 for MPPCA with Rician noise.

The reconstructed images with both NORDIC and MPPCA shown in **Figure 3A** (middle and lower row), all appear qualitatively similar to the reference image (Fig.3A.i), with the MPPCA applied on data with Rician noise showing evidence of the Rician noise-floor. The *q*-space contrast images (**Figure 3B**) show that for the patch sizes 5^3^ with NORDIC and for both MPPCA Gaussian and MPPCA Rician, there is significant deviation from the reference image (FIG.3B.i), especially in the cerebellum area. For **NORDIC**-processing with patch sizes of 5^3^, 7^3^, 11^3^, 15^3^, and 19^3^ the structural similarity index (SSIM) (Wang et al., 2004) over all slices and diffusion direction is shown in **Figure 3C**. The SSIM is highest for a patch size of 11^3^ and for the subsequent application of **NORDIC** a patch size of 11^3^ is used unless specified otherwise.

### In-vivo Imaging

#### Impact of phase-stabilization and patch averaging

For LLR techniques applied on complex-valued data, the necessity and impact of phase-stabilization is shown along with the gains from patch averaging in **Figure 4** for a diffusion weighted image with b=3000 s/mm^2^ from the 0.9mm acquisition and processed with NORDIC using an 11^3^ patch. The images shown are reformatted oblique coronal slices extracted from the 3D volume, acquired as oblique axial, so that the vertical axis now corresponds to the slice direction. Without phase-stabilization, the diffusion phase has a high frequency fluctuation along the slice direction. This limits the efficacy of the LLR representation model, not allowing for an effectively low-rank representation. In the reconstruction without patch averaging or phase-stabilization, the underlying anatomy is not recovered efficiently, and horizontal stripe-like intensity variations are observed. These stripe-like intensity variations are more easily observed in the reconstruction (2^nd^ from left) obtained without phase-stabilization and with patch averaging. With patch averaging and phase-stabilization, the underlying anatomy is clearly observed both peripheral and medial in the coronal view. Note that with averaging, for a patch size of 11^3^, each voxel is averaged from 11^3^ reconstructions, further improving final image quality. Unless otherwise noted complete patch averaging is used along with phase-stabilization.

**Figure 4.**
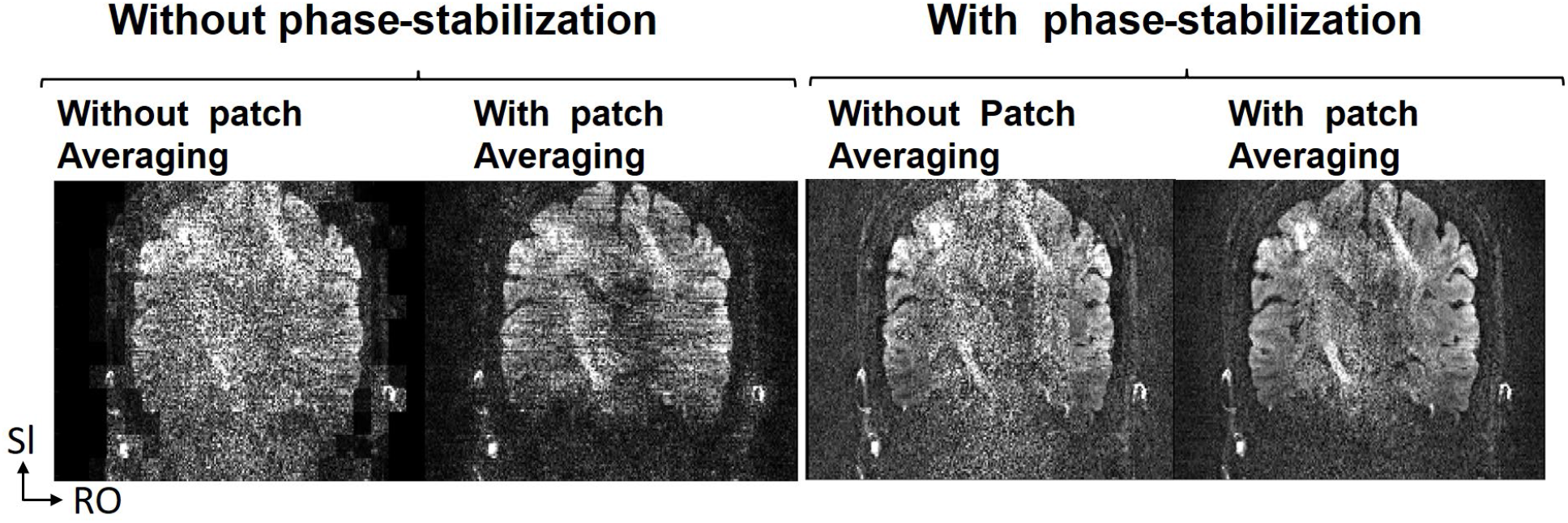
Effect of phase-stabilization and patch averaging for the NORDIC processing of complex data using an 11^3^ patch. Reformatting to oblique coronal slices of the 0.9mm isotropic data, acquired with oblique axial acquisition, are shown. The left figures are without phase-removal of the slice-specific smooth diffusion phase, and the right figures are with removal of the diffusion phase. For both approaches, the impact of patch-averaging with all 11^3^ patches are shown.

#### Performance of NORDIC at Different Resolutions

**Figure 5** illustrates an axial slice of the FA map obtained after dMRI processing (top row), and the image of the corresponding slice from the volume with b=3000 s/mm^2^ weighting (lower row), for the three different resolutions and a single subject. The FA maps and the corresponding slice image with b=3000 s/mm^2^ are presented for the standard- and the **NORDIC**-processed data, for the different resolutions. The signal scaling for the bottom row is adjusted for each resolution, since SNR varies with resolution.

**Figure 5.**
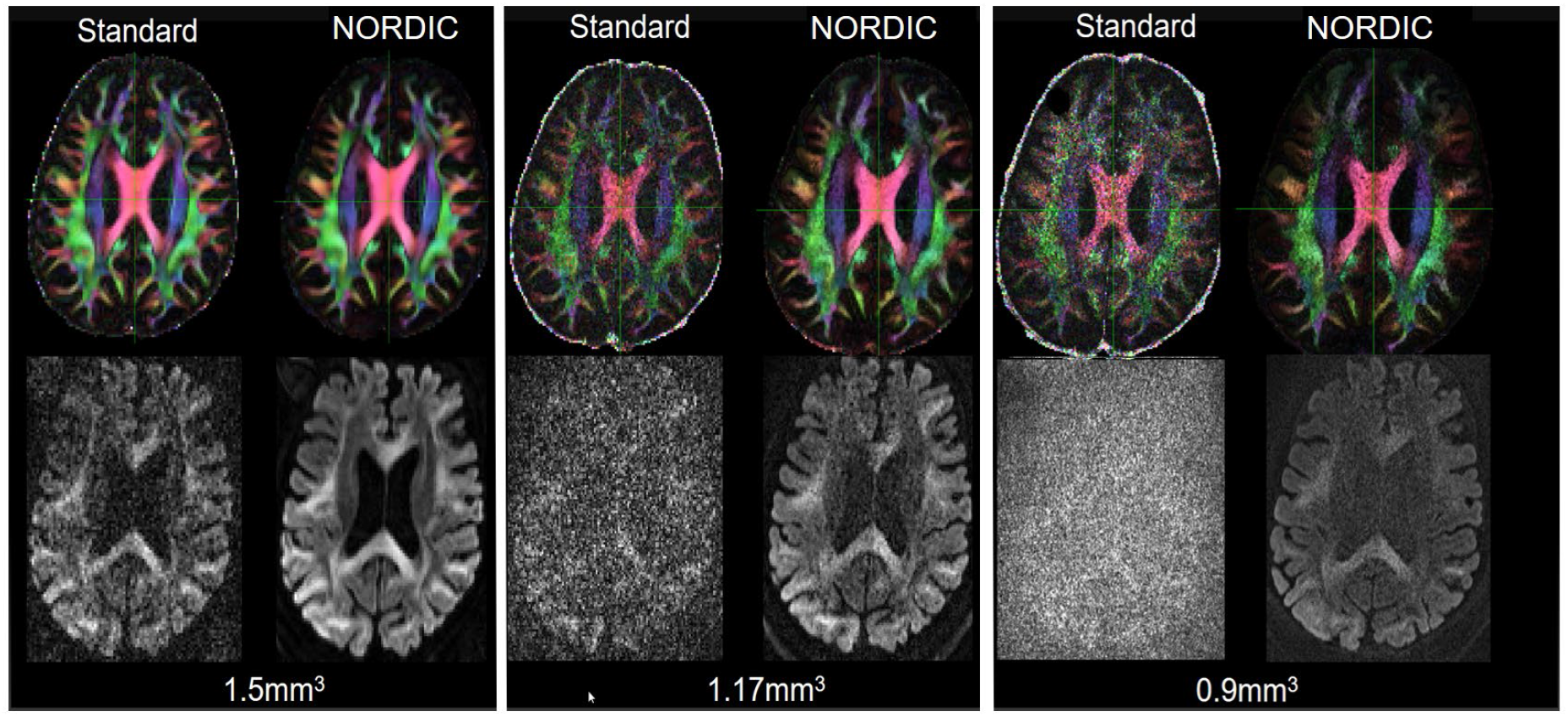
Top row shows the effect of NORDIC across different resolutions on FA maps for a single slice. Bottom row presents corresponding diffusion weighted image (b=3000 s/mm^2^) for the same slice. The FA maps are obtained after EDDY processing, and the images in the bottom row are before EDDY processing. From left to right, in groupings of 4, the three resolutions of 1.5mm, 1.17mm and 0.9 mm are shown. For each grouping, the images with the standard processing are shown adjacent to the images with the NORDIC processing. Supplemental Figure S3, illustrates the same slices and also includes reconstruction by MPPCA for comparison.

The spatial tissue contrast is similar in the **NORDIC**-processed images of the slice illustrated in the lower row in **Figure 5** for the 1.5mm or 1.17mm resolutions. In the 0.9mm resolution image, the noise over the lateral ventricles is not suppressed as much as it is in the 1.5mm or 1.17mm resolutions. Some differences in the overall anatomical features between the different resolutions can be noted and are due to slight differences in slice position and thickness. The standard and **NORDIC**-constructed FA maps for the 1.5mm resolution are visually identical, which also illustrates the high SNR quality of the HCP lifespan protocol. For the 1.17mm and 0.9mm resolutions, the FA maps are noisier compared to the 1.5 mm data, but have similar features with the lower resolution FA map. The FA in this context is also an adequate visual measure for the resolution that is maintained with **NORDIC**, as there is no evident blurring, consistent with the analysis from (Veraart et al., 2016), which compared MPPCA with adaptive non-local means and second order total generalized variation. Corresponding results for MPPCA are shown in Supplemental Figure S3.

#### SNR Advantage of NORDIC: Comparison with Data Averaging

For the 3 subjects with repeated acquisitions, the effect of **NORDIC** after EDDY correction is shown in **Figure 6** for an axial slice in each subject. In this case, each single “repetition” refers to the pair of separate acquisitions with reversed phase encoding which is used for EPI corrections. TOPUP/EDDY, combines data with opposite phase-encoding directions, improving the SNR by approximately 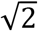 compared to a true single acquisition.

**Figure 6.**
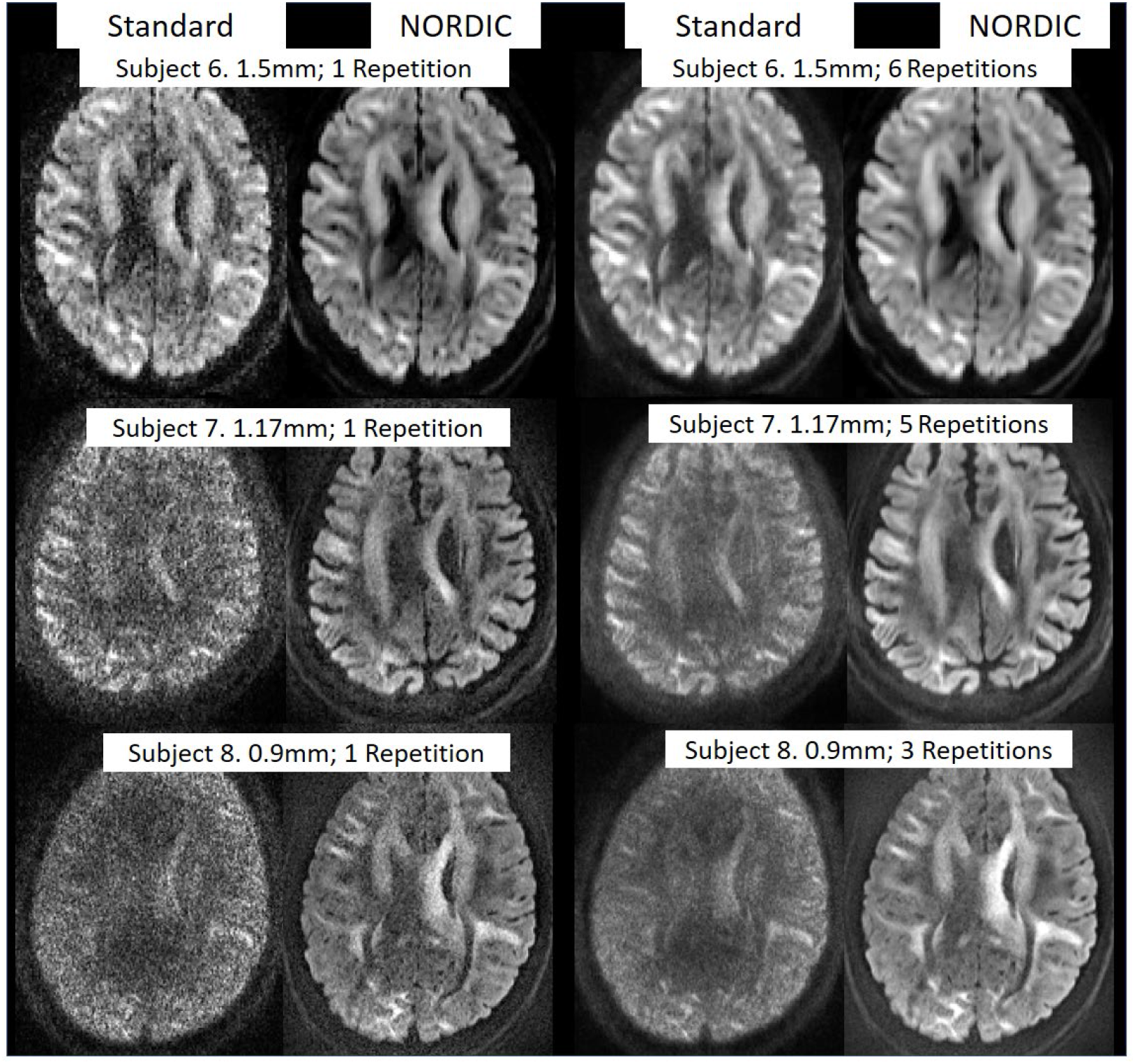
Comparison of NORDIC processing with averaging of repetitive acquisitions to increase SNR. The left two columns are for a single acquisition across 3 different resolutions and the right two columns are for the averaging of the repetitive acquisitions. In each case, EDDY processing was applied. Supplemental Figure S4 illustrates the same data and also includes reconstruction by MPPCA for comparison.

For the 1.5 mm resolution data, the **NORDIC**-processed single repetition data visually shows reduced noise in comparison with the standard reconstruction, and is similar to the standard reconstruction obtained by the averaging of 6 repetitions. For the 1.17mm resolution, the single repetition processed with NORDIC (2^nd^ column) displays improved SNR on visual inspection compared to the average of the 5 standard reconstructions (3^rd^ column), especially for the deeper brain regions which have intrinsically lower SNR. We note that the single repetition also has similar anatomical features to the 5 repetitions averaged after processing both with **NORDIC,** and that in this case, the average of the 5 repetitions processed with **NORDIC** has a higher visual SNR, as would be expected, compared with the single repetition data reconstructed with **NORDIC**. For the 0.9mm resolution, the **NORDIC**-processed single repetition and 3 repetition data has clearly defined anatomical features, which are barely perceptible in the standard reconstruction images, even when 3 acquisitions are averaged to improve SNR. Corresponding results for MPPCA are shown in Supplemental Figure S3, which shows that the effect of the Rician noise reduces the contrast of the individual denoised images.

#### Quantitative evaluation of patch size for NORDIC

**Figure 7** illustrates quantitatively an dMRI data from a single subject, the performance of denoising with **NORDIC** applied with different patch sizes; the metrics shown are the 2^nd^ and 3^rd^ *fiber detection rate* (percentage of voxels within a VOI with two- and three-way fiber crossings) and *gain in fiber orientation accuracy* (i.e. the decrease in the angular dispersion (uncertainty) in determining the fiber orientations) in the two brain regions well-known to have second and third fiber crossings, the PCR and SLF. The rightmost column in **Figure 7** shows the atlas-based definition of the SLF and PCR used for quantification of crossing fibers; all voxels within the regions shown by the green and red colors were used for the analysis.

**Figure 7.**
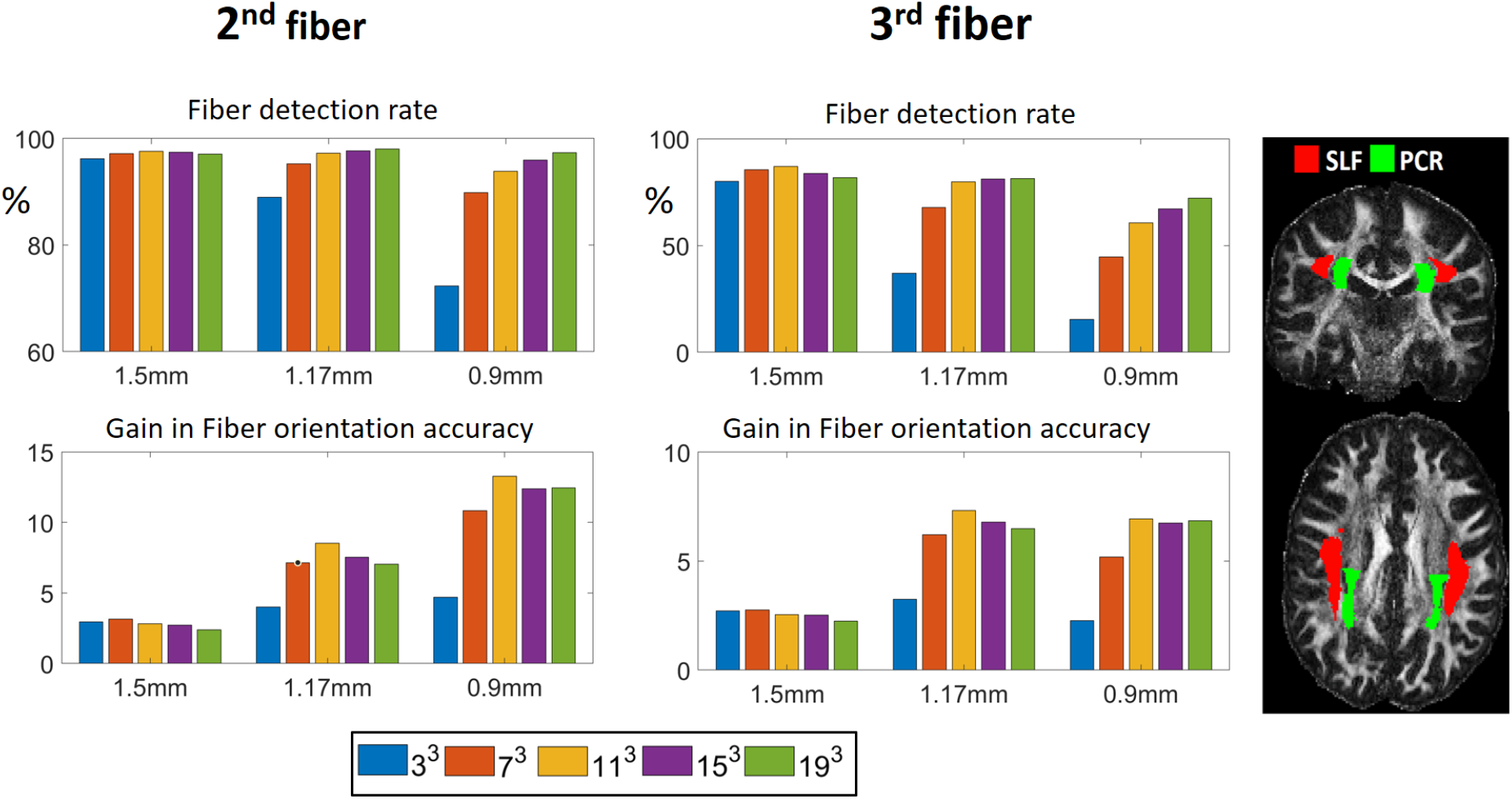
Effect of patch size in **NORDIC** processed dMRI data on fiber detection rate and accuracy for the three different resolutions. The 5 different patch sizes compared are with *n^3^* for *n* = {3,7,11,15,19}. The top row, shows the detection rate of voxels which supports a second and third fiber, and the bottom row shows the gain in fiber orientation accuracy after **NORDIC** relative to the standard processing as a ratio between the dispersion determined for the standard and NORDIC processing. For MPPCA, the fiber detection rates were [94%, 77% and 38%] for the 2^nd^ fiber, and [73%, 31% and 8%] for the 3^rd^ fiber, with gains in Fiber orientation accuracy of [2.2, 3.4, and 3.0] for the 2^nd^ fiber and [2.2, 3.3 and 1.9] for the third fiber.

In the top row, the fiber detection rate shows that for the 1.5mm resolution data, a high percentage of voxels in these VOIs supports both a second and third fiber; for the 1.5mm data set, there is minimal impact of the patch size either on the detection rate (top row, **Fig. 7**) or the gain in fiber orientation accuracy (bottom row, **Fig. 7)** as the ratio between the dispersion determined for the standard and NORDIC processing. The impact of the patch size is much more pronounced for the higher resolution data, which have intrinsically lower SNR. For 1.17mm resolution, the detection rate of voxels supporting a second and third fiber increases with the patch size and plateaus at the 11^3^; the gain in fiber orientation accuracy increases correspondingly and is largest for the 11^3^ patch size and then degrades for increasing patch sizes without a corresponding increase in the number of fibers being detected. Similarly, for the 0.9mm resolution data, the gain in fiber orientation accuracy is highest for the 11^3^ patch size; in this case, however, the detection rate of voxels supporting a second and third fiber increases monotonically with increasing patch size, albeit, the reliability of those added fibers is low. This further suggests that, for a dMRI series with 99 q-space samples a patch size of 11^3^ achieves the best trade-off across resolutions.

#### Fiber Quantification Performance of NORDIC

The trends of *fiber orientation dispersion* reflecting the uncertainty in the fiber orientation estimation and the reduction in this dispersion as a *gain in fiber orientation accuracy* with **NORDIC** are shown in **Figure 8,** for all dMRI data acquired (i.e. for the 5 subjects with a single acquisition at the three resolutions, and for the 3 subjects with repeated acquisitions at a single resolution). For the 5-subject data, the height of the bars in each plot represents the mean uncertainty and the mean gain in fiber orientation accuracy, calculated for the single (i.e. 1 repetition) acquisitions obtained from the 5 individuals; in this case, the error bars represent the standard deviation across subjects. These are plotted adjacent to single subject data (a different subject for each resolution), but acquired multiple times, with the height of the bars representing the mean of the repeated single acquisitions and the error bars representing the deviation among the different single acquisitions acquired in different sessions.

**Figure 8.**
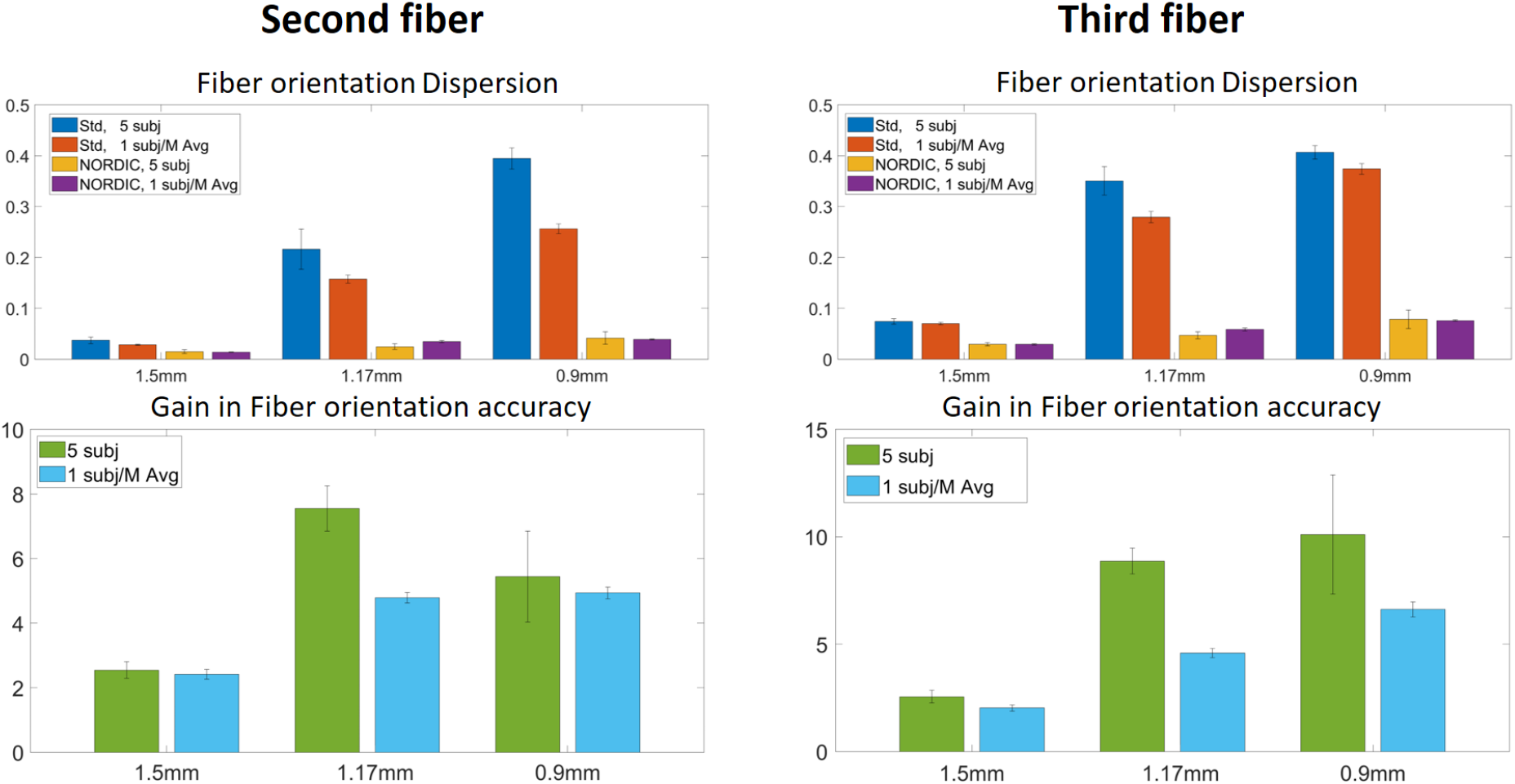
Quantitative metric in brain regions SLF and PCR for standard and NORDIC processed data for the 5 subjects scanned at different resolutions and the 3 subjects each scanned at a single resolution with multiple repetitions. The top row illustrates fiber orientation dispersion (reflecting the uncertainty in the fiber orientation estimation) for voxels within a VOI supporting a second fiber (left), and supporting a third fiber (right); for this metric, lower height of the bar indicates better performance (lower uncertainty). The *gain in fiber orientation accuracy* (i.e. a decrease in dispersion reported as the ratio of the dispersions calculated with standard to that calculated with **NORDIC** processing) is shown in the lower row for the voxels supporting second (left) a third fiber (right); for this metric, the higher bar indicates better performance for NORDIC. The rightmost column in Figure 7 shows the segmentation of the SLF and PCR used for quantification of crossing fibers. The error bars for the *multi-resolution single repetition* data represents the variability between subjects, and the error bars for the *single-resolution multiple repetitions* shows the variability within subjects but over different acquisitions. In case of MPPCA for the *single-resolution multiple repetitions*, gains in Fiber orientation accuracy were [2.2, 3.4 and 3.0] for the 2^nd^ fiber for the 1.5, 1.17 and 0.9 mm resolution data, respectively; the corresponding numbers were [2.2, 3.3 and 1.9] for the third fiber with MPPCA processing.

**NORDIC** processing leads to major improvements in these diffusion metrics, as shown in **Figure 8** both for the single and multiple acquisition(s) data obtained at each spatial resolution. For the single resolution and multiple acquisitions, the variability in gain in fiber orientation accuracy is more consistent than the variability in gain across subjects. Moreover, the absolute gain in accuracy (lower row **Fig.8**) between the two groups is similar for the 1.5mm and 0.9mm resolution. For the 1.17mm the subject with repeated acquisitions had a lower uncertainty in the fiber dispersion before **NORDIC** processing, and the gain in fiber orientation accuracy in this case was not as large as for the group with multiple resolutions and single repetition. This is consistent with results shown in **Figure 6**: After **NORDIC** processing, the uncertainty in the single acquisition and the multiple acquisition groups became similar.

In **Figure 9**, the fiber detection rate within a VOI in the brain regions SLF and PCR is plotted against the fiber orientation dispersion from the multi-resolution single repetition data and from the single resolution multiple repetitions data. The plots shown in the top row are for voxels supporting second fibers, and those in the bottom row are for voxels supporting a third fibers.

**Figure 9.**
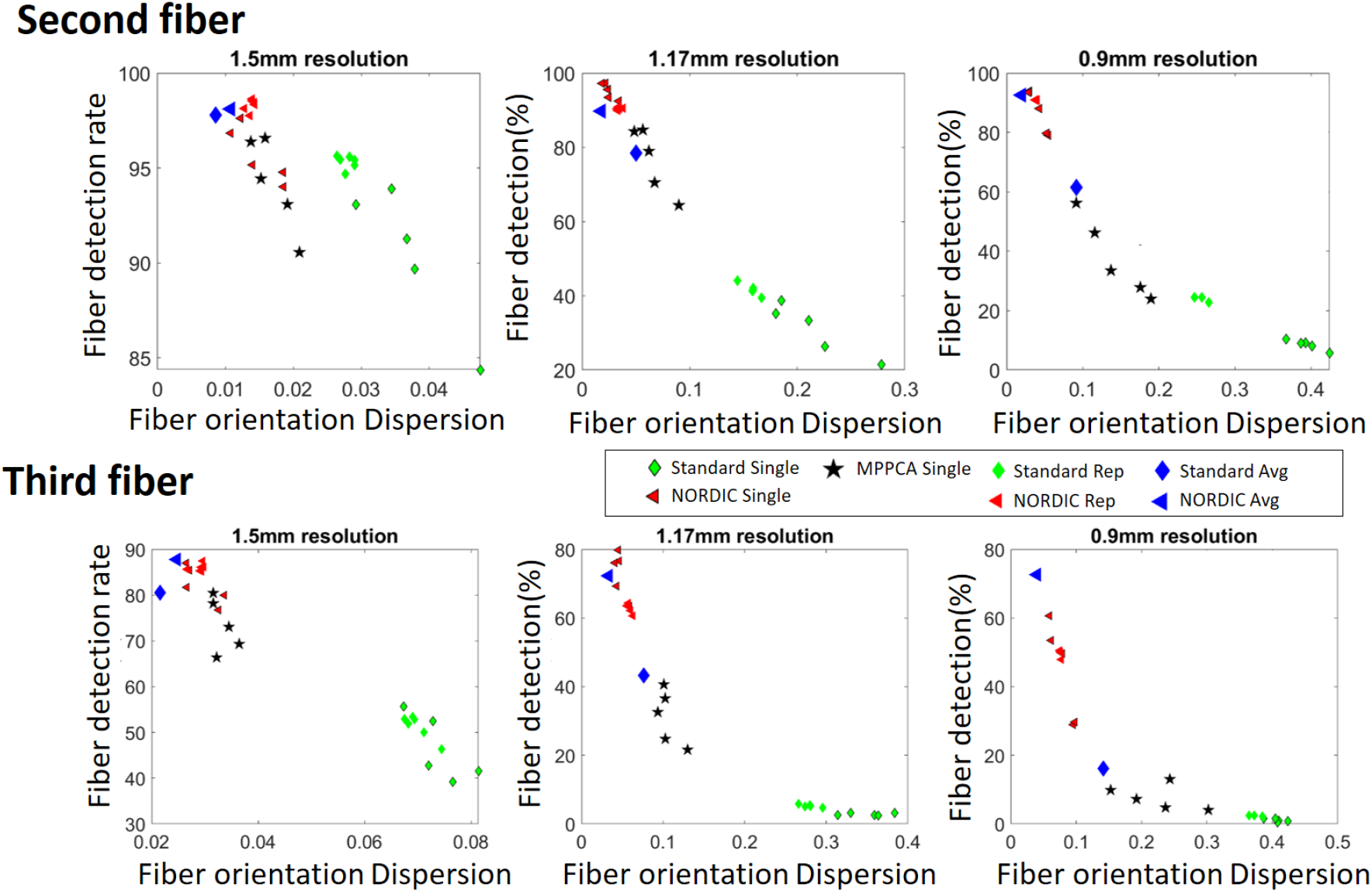
Scatter plot of the *detection rate* of voxels with second (top row) and third (bottom row) fibers against the fiber orientation dispersion, reflecting the uncertainty in the fiber orientation estimate for the fibers in the brain regions PCR and SLF after bedpost processing. The VOI is determined from the subject independent JHU-ICBM atlas, and resampled to the data-space for each subject. The vertical axis (detection rate) is expressed as % of voxels in the VOI that contain two or three fiber crossings (second and third fibers, respectively). For *multi-resolution single repetition* the standard Single, **NORDIC** Single and MPPCA Single processing are shown, and for *single-resolution multiple repetition* the standard Rep and **NORDIC** Rep processing are shown. The Standard Avg and **NORDIC** Avg are extracted from dMRI data after averaging of multiple repetitions obtained at a single resolution, where each acquisition is independently corrected with EDDY, then jointly motion corrected and averaged before processing with bedpostX.

##### Multi-resolution single repetition data

In each plot, the green diamonds with black outline designate the values for the standard reconstruction, the red triangles with black outline designate the values for NORDIC processed data and the black stars designate the values for MPPCA. With the application of **NORDIC,** the change in dispersion of the second fiber for the 1.5mm, 1.17mm and 0.9mm data, reflect a gain in fiber orientation accuracy (a decrease in dispersion) of a factor 2.6, 8.9 and 10.1, respectively, for the second fiber, and 2.5, 7.6 and 5.4 for the third fiber (green bars in **Figure 8**). For the 1.5mm, 1.17mm and 0.9mm with **NORDIC** the detection rate of voxels with second fibers was about the same at 96%, 95% and 87% of the atlas based VOI, while the detection rate of voxels with third fibers was 82%, 73% and 45%. This is not surprising, as detection of third fibers is more sensitive to higher noise levels in the higher resolution data.

Using **NORDIC** processing, resulted in gains in the metrics plotted in **Figure 9**, fiber detection and orientation accuracy, compared to the MPPCA technique; in this two-dimensional plot, the **NORDIC** points appear higher in detection rate axis and lower (towards the left) on the fiber orientation dispersion axes compared to the MPPCA processed data, particularly for the lower SNR 1.17 and 0.9 mm data. Looking at the fiber orientation dispersion metric alone, with NORDIC denoising relative to MPPCA, the gain in orientation dispersion for the average of 2^nd^ and 3^rd^ fiber is higher with NORDIC by a factor of 1.15, 2.45 and 3.3 for the 1.5mm, 1.17mm and 0.9mm data, respectively.

##### Single-resolution multiple repetition data

In each plot, the green diamonds and red triangles without a black outline are the standard and **NORDIC** processing of the single repetition data, respectively from the pool of data where a single individual was scanned multiple times for a given resolution. With the application of **NORDIC,** the change in dispersion of the second fiber for the 1.5mm, 1.17mm and 0.9mm data, reflect a gain in fiber orientation accuracy (a decrease in dispersion) of a factor 2.0, 4.6 and 6.6, respectively, for the second fiber, and 2.4, 4.8 and 4.9 for the third fiber (blue bars in **Figure 8**).

The blue diamonds and triangles are the average of the repeated acquisitions processed with standard and **NORDIC** processing, respectively, where after reconstruction each acquisition is independently corrected with EDDY, then jointly motion corrected and averaged before processing with bedpostX.

For the 3 subjects with multiple repetitions, the dispersion of the second fiber for the SLF and the PCR were calculated for each individual series separately and after averaging the multiple repetitions of the dMRI data. For the standard data *without* **NORDIC**, the average gain in uncertainty was 3.2, 3.1 and 2.8 for the 1.5mm (with 6 averages), 1.17mm (with 5 averages) and 0.9mm (with 3 averages), respectively. The gains are slightly higher than the direct gains in image SNR of 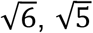, and 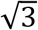 respectively, and supports using the gain in fiber orientation accuracy as a proxy of SNR.

After processing each series with **NORDIC**, the fiber orientation dispersion of the second fiber of the *averaged series* for the 1.5mm resolution changed by factor 1.28 relative to each individual series. For the 1.17mm resolution, the fiber orientation dispersion changed by 1.94 relative to each individual series, and for the 0.9mm resolution, the fiber orientation dispersion changed by 1.97 relative to each individual series.

The smaller change noted for the 1.5mm indicate that for a single series, after **NORDIC**, the dMRI model is not able to better describe the underlying properties, since the standard data already has good SNR, whereas the SNR of the 1.17mm and 0.9mm individual scans is more substantially improved by **NORDIC**.

#### Impact of NORDIC for Whole Brain Tractography

In order to demonstrate the whole brain effect of improved detection of crossing fibers in the **NORDIC** processed data, the connectivity of the entire subject specific PCR from **Figure 7** (both left and right hemisphere) with the rest of the brain was investigated qualitatively using probabilistic tractography (‘probtrackx’)(Behrens et al., 2007). The connection strength, defined as the number of streamlines (extracted by the tractography algorithm) passing through each brain voxel, and connecting the PCR to the rest of the brain, is used to represent the connectivity (Behrens et al., 2007). **Figure 10a** shows sagittal, coronal, and axial views of the connectivity distribution for the 0.9mm data from the probabilistic tractography for standard processing of a single acquisition (left), the average of the repeated acquisitions (right), and the **NORDIC** processed single acquisition (middle).

**Figure 10.**
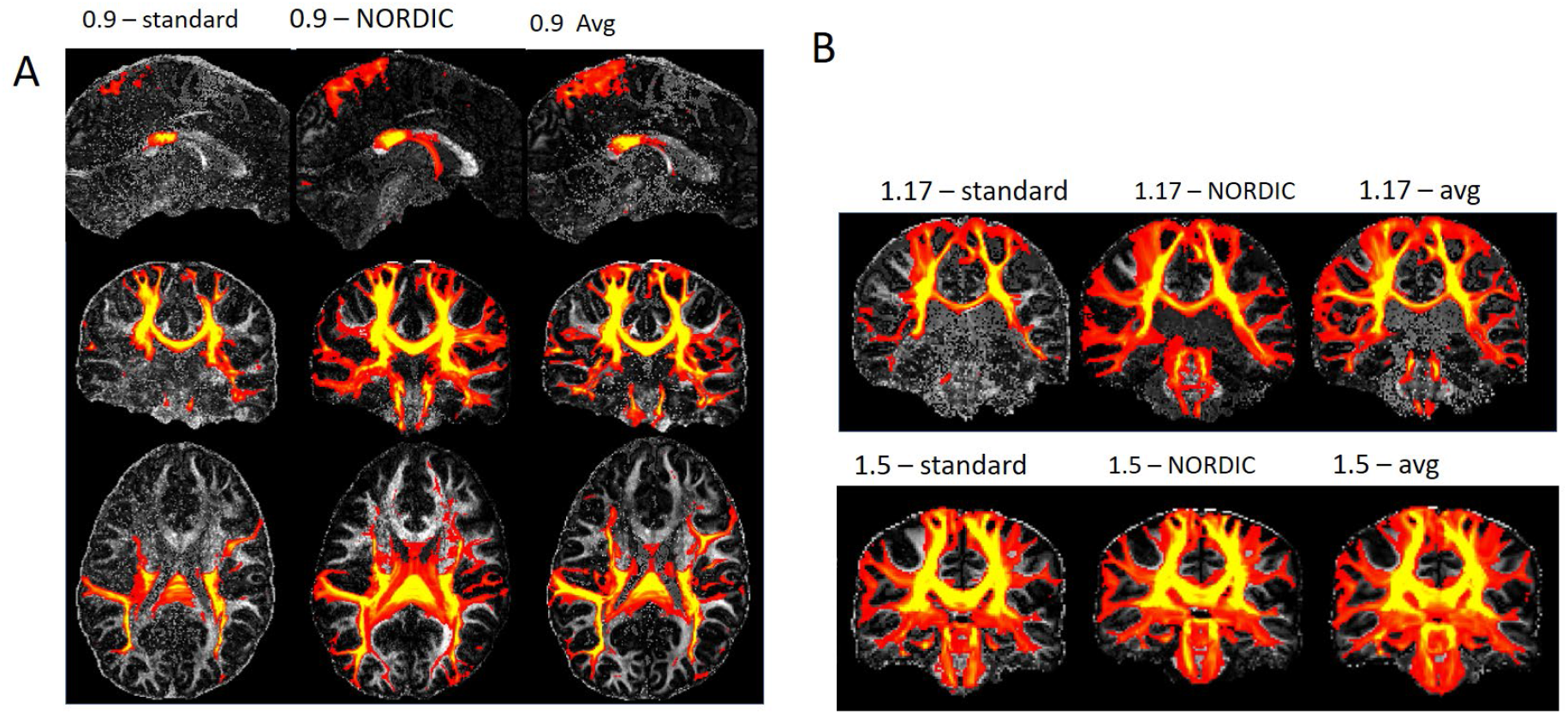
Comparison of connectivity distributions from the probabilistic tractography results for 0.9mm data (A), and 1.17mm ((B) upper panel) and 1.5mm ((B) lower panel) data representing connectivity of the entire subject specific PCR.

The regions covering the main white matter tracts near the seeds, in both left and right PCRs, show high connection strength in the standard data, and with the NORDIC-processed data. These connectivity distributions are expanded along the tracts with increased symmetry, for example along the corticopontine and corticospinal tracts. The improved probabilistic tractography of the fiber bundles passing through the PCR connecting it better to the rest of the brain, which is a result of the improved detection of second and third crossing fibers along with reduced orientation uncertainty, is evident in the single acquisition **NORDIC** data, compared to the standard data. In addition, the figure also shows improved connectivity in single acquisition **NORDIC** data compared to the standard average data, especially around the brainstem. **Figure 10b** shows sample sagittal views of the connectivity distribution from the probabilistic tractography results for 1.17mm (upper panel) and 1.5mm (lower panel) data, for single acquisition (left), single acquisition **NORDIC** (middle), and average of multiple acquisitions (right). For the 1.7mm and 1.5mm acquisitions, the improved connectivity profile from the cortex through the PCR to the pons can be seen, and the **NORDIC**-processed 1.17mm data exhibits similar sensitivity for the connectivity as in the standard 1.5mm data. These observations can be partly explained by the improved sensitivity to crossing fibers, and performance of the **NORDIC**-processed data, as demonstrated in the quantitative comparisons in **Figure 9**. In these experiments, we showed that the orientation dispersion (the proxy for SNR) of the second and third fiber, after the use of **NORDIC,** is less than 0.1 across all resolutions, the detection rate of voxels with a second fiber is in excess of 80% of the atlas based voxels, and the detection rate of voxels with a third fiber after **NORDIC** for the higher resolutions is higher than the standard processing for the lowest resolution. Corresponding results for MPPCA are shown in Supplemental Figure S5, from which improved connection strength with NORDIC compared to MPPCA processing, is evident in the corpus callosum area for high resolution acquisitions. At 1.5mm, the difference between MPPCA and NORDIC processed data is less compared to that at higher resolutions, mainly because the number of second fibers estimated at 1.5mm is somewhat similar between these methods, but the improvement in connection strength (the width of the connections shown in yellow) is visible which is due to the lower orientation dispersion and higher number of third fibers resolved in the NORDIC processing.

The effect of NORDIC is demonstrated on tractography streamlines with two high resolution data in Figure 11. Figure 11A, B show the streamlines constructed before and after NORDIC denoising, from the same 0.9mm data which was used for all the previous analysis presented in Figures 1 through 10. The relatively poor SNR of this acquisition results in tractography streamlines that are clearly problematic on visual inspection, most obviously evidenced by the discontinuities in the normally prominent corticospinal tracts and the almost randomly oriented appearance of the streamlines near the cortical surface. The use of NORDIC fixes these problems. This improvement in the tractography streamlines are fully consistent with the probabilistic tractography results given in Figure 10. A more dramatic improvement is shown in Fig.11C and D, using a 0.7mm isotropic whole brain dMRI data. In this 0.7mm data, without denoising the tractography completely fails except in the corpus callosum. After denoising the commonly seen tracks are now clearly detectable.

**Figure 11.**
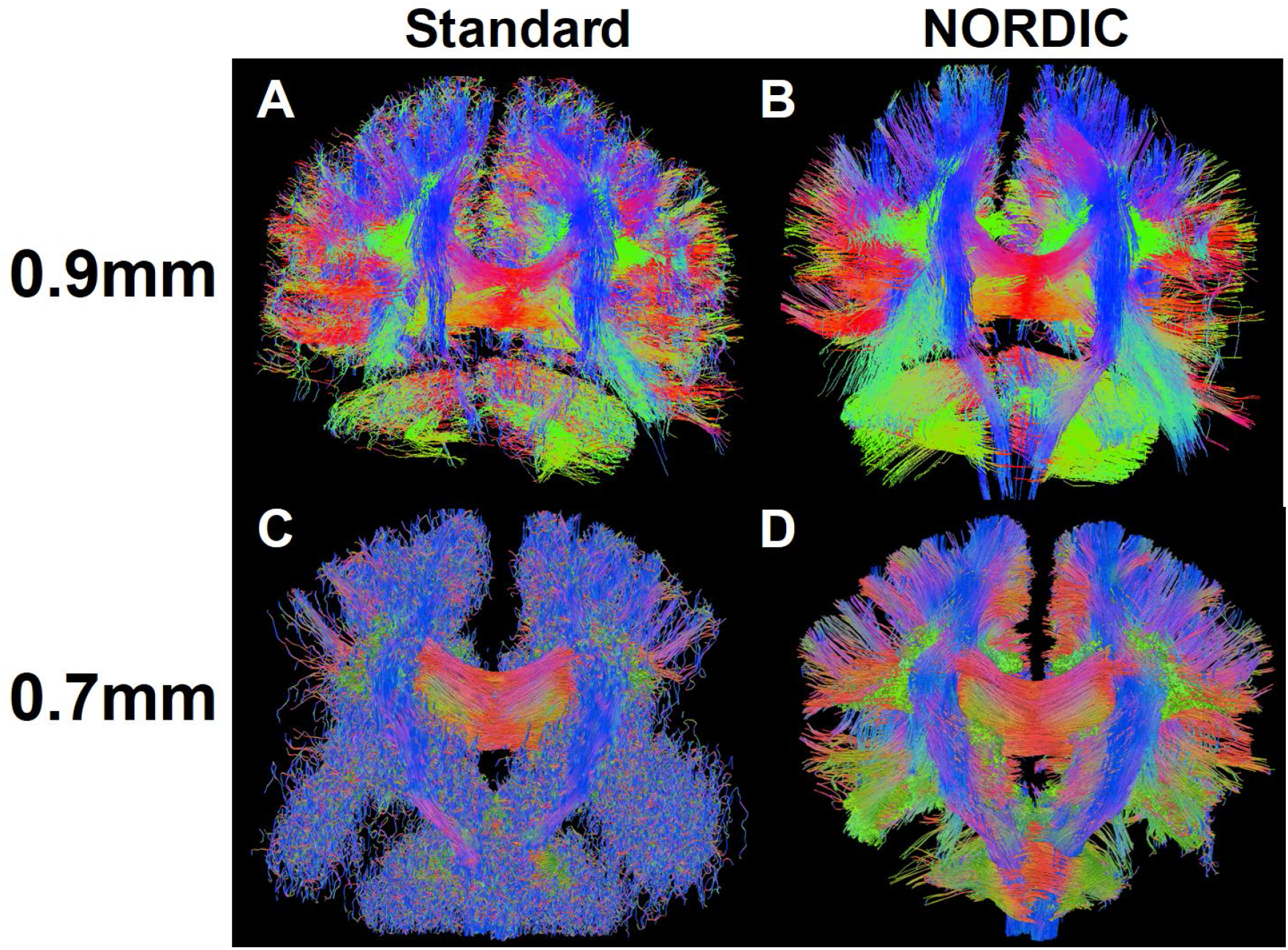
Comparison of tractography streamlines without and with NORDIC processing. The top row shows a comparison for the 0.9mm isotropic resolution data analyzed in Figure 5 to Figure 10 using a single repetition with standard processing (A), and with NORDIC processing (B). The bottom row, likewise shows a comparison for the 0.7mm isotropic resolution data with standard processing (C) and with NORDIC processing (D). This improvement in the tractography streamlines are fully consistent with the probabilistic tractography results given in Figure 10. The 0.7 mm data set has different acquisition parameters and as such cannot be directly compared to the 0.9 mm data *per se;* it is included here only to show a more dramatic improvement possible with lower SNR data.

## Discussion

In this study on the impact of locally low rank constrained processing for dMRI, we have proposed a pre-processing pipeline and a parameter-free thresholding technique based on the known properties of Gaussian thermal noise by selecting the largest singular value of i.i.d. noise. We jointly refer to this thresholding technique and pre-processing pipeline as the NOise Reduction with DIstribution Corrected (NORDIC) PCA. While the use of the spectrum of random noise is reminiscent of the MPPCA method, there are several key differences. MPPCA is typically implemented to work on DICOM images, which do not generally fit the assumptions in the Marchenko–Pastur law and its theoretical bounds. In our approach, this issue of not having i.i.d. zero-mean noise components was resolved by using complex-valued processing and spatial noise correction to match with the Marchenko–Pastur law. The g-factor for the k-space reconstruction method is determined from the reconstruction parameters and captures precisely regions with rapid changes in thermal noise. The g-factor based spatial noise correction enables a consistent use of thresholding across regions with otherwise rapidly changing spatially non-uniform thermal noise levels. In dealing with complex data, we further proposed a phase-stabilization approach to promote the low rank nature of patches by explicitly reducing the fluctuations in the phase of the MR signal subsequent to diffusion encoding during q-space sampling; this phase is not used for dMRI analysis, and removal of its variations does not impact the information inferred from the diffusion encoding. Additionally, in MPPCA, the threshold for denoising is determined from the data itself by estimating the number of components from a singular value decomposition that can be summed while still being within the limits of the asymptotic spectral bounds of random matrices. This favors looking for sharp transitions in the spectra, reflecting an underlying low-rank signal; even though visually evident, this is not a guaranteed criteria and when missed removes too many signal components as well as noise. In contrast, in our approach, the largest singular value of the known thermal noise is selected, which is equivalent to removing all components that cannot be distinguished from random Gaussian noise.

MPPCA as a default implementation in (Veraart et al., 2016) and in dwidenoise in MRTricks for the application of the Marchenko-Pastur based technique for DICOM (magnitude) images, uses a 5^3^ patch for the Casorati matrix, with a *fast* mode where the patches are not overlapping and a *full* mode where they overlap. The implementations can also be applied to complex images, but without additional modifications that ensure a spatially uniform noise distribution; consequently, the performance is inferior to the magnitude implementation. The determination of significant signal components for MPPCA is elegant in that, for local patches, it calculates a threshold based on the tail of the spectrum from the analytic decay properties of random noise, but it is conceptually challenging to know what threshold was determined and if the data has properties that could be better exploited.

For **NORDIC** we demonstrate that a patch size of 11^3^ is better for a series with 99 volumes in maintaining the correct contrast across q-space acquisitions, corresponding to a ratio of approximately 11:1 of the M×N patch sizes used for NORDIC processing. While larger patch sizes are also beneficial for reaching the asymptotic conditions for the Marchenko-Pastur, larger patch sizes are not good for LLR techniques since the large patches become highly heterogeneous with respect to tissues of different kinds and, as such will deviate from being low-rank and will make the determination of the threshold more difficult. The spread in the square of the spectra of the noise increases only as 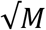, and for MPPCA the trade-off in patch size was found to be 5^3^ such that the spectrum has a large slope, which more quickly provides a contrast when the joint singular value spectrum has both a signal and noise contribution. For larger patches, small benefits relative to the computational challenge was also found (Ma et al., 2020). Although it should be noted that for large kernels such as 11^3^ using patch overlap of just ½ the FOV creates virtually identical reconstructions to the maximal overlapping reconstruction, while also reducing the computation 50 fold and making it comparable with 5^3^ kernels in terms of computation time. With this, a simple matlab implementation of 11^3^ patches is faster than the 5^3^ for both **NORDIC** and MPPCA.

For **NORDIC**, the absolute noise-level in image space is assumed to be known which is not normally tracked since most MRI images are not reconstructed in absolute SNR units as proposed in (Kellman and McVeigh, 2005), but are typically coarsely scaled to maximize the precisions in the data-type used. The system calibrations performed prior to each acquisition has the information about the k-space noise-level. In our implementation, a synthetic signal based on the i.i.d. noise estimate initially was used to artificially create an extra volume in the series. The noise-level in image space after removal of the g-factor noise can also be estimated from a region without signal, or alternatively an acquisition without RF excitation can be added. All of these approaches provide the same information needed for selecting the **NORDIC** threshold. Likewise one can hypothesize that an absolute shift-invariant noise-level can be obtained with the MPPCA as a mean estimate of each locally estimated noise-level with (eq.2), or as proposed for MRI by Foi et al (Foi, 2011) using the variance stabilizing transform.

The denoising with either MPPCA on magnitude images or NORDIC on complex images recovers the underlying image when the SNR is high enough (Figure 5 and Supplemental Figure S3); however, as the SNR decreases the impact of the non-zero iid thermal noise becomes more apparent for the MPCPA processed data as revealed in a loss of q-space contrast and residual high-spatial frequency modulations. This is reflected less in an average measure such as the FA map, but quantitatively shown for the fiber orientation dispersion (Figure 9 and Supplemental Figure S5). The recently proposed VST algorithm (Ma et al., 2020) confronts the difficulty of working with magnitude data using a two-step approach to resolve the Rician and the spatially varying noise and then uses MPPCA for denoising. Although this approach provides improvements over simply working with magnitude data with Rician noise distribution, it also introduces spatial smoothing which the NORDIC approach does not. One such example is shown in Supplemental Figure S6 for both 1.17mm and 0.9mm isotropic resolution with b=3000s/mm^2^. Using AFNI tools(Cox, 1996) for Gaussian blurring through estimation of auto-correlation function among voxels, for VST an increased blurring of 50-100% for the 1.17mm and 0.9mm isotropic resolutions was measured. Such blurring is in general undesirable especially when high resolutions are intentionally targeted for improved tractography Nevertheless, for data that exists only in DICOM format, as they do for example in many large databases generated to date and still being collected in many laboratories, VST is a better option then applying MPPCA directly as exhaustively demonstrated by Ma et al (Ma et al., 2020). However, if complex valued data can be saved, as we are sure will be going forward, NORDIC is the preferred image reconstruction method.

The application of LLR assumes a low-rank signal, and sporadic or random motion increases the underlying rank of the image series, since the low-rank model needs to encode both the bio-physical signal properties and the motion model. As such, motion corrected data can be better corrected for thermal noise fluctuations. For dMRI, eddy-currents specific to the diffusion encoding direction and magnitude introduces volume specific distortions, which increases the volume-to-volume variability. While correcting for these fluctuations with post processing algorithms like EDDY before noise-removal increases the volume-to-volume anatomical consistency, it also changes the noise-properties. Furthermore, since EDDY is known to correct for higher order effects, such as slice-dependent signal dropout from motion and volume specific susceptibility distortion correction it warrants further investigation what effects are obtainable if the LLR and EDDY corrections are switched or integrated jointly.

The known image reconstruction parameters of sensitivity profiles, determined with ESPIRIT, for the SENSE-1 reconstruction, and the slice-GRAPPA convolutional kernels, determined jointly for MB×R acceleration, are used through the pipeline to provide the desired noise-properties of i.i.d. Gaussian without introducing additional estimations. This furthermore provides as a fast and mathematically exact calculation of the known parameters as auxiliary information that should be beneficial for general quantification of parameter mapping since it facilitate exact quantification of spatial SNR for MB×R acquisitions. The use of NORDIC preserves subtle consistent features in the images, which may include Nyquist ghosting and residual slice-aliasing.

Under controlled conditions, and using metrics from the HCP for dMRI information content evaluation, acquisitions with 1.5mm, 1.17mm and 0.9mm were used. The 1.5mm protocol was aligned with the lifespan protocol and the sequence parameters for the 1.17 and 0.9mm protocols where selected such that they weighted towards shorter TR, *versus* higher image SNR. This required aggressive use of combined slice and phase-encoding undersampling. The 1.17mm had about 5-fold less SNR than the 1.5mm (2-fold reduced volume, 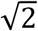 for phase-encoding undersampling, g-factor with acceleration of MB×R=5×2 versus MB×R=4×1, and 20% higher signal from shortened TE from phase-encoding undersampling); this SNR difference also corresponded to mean fiber orientation dispersion in the PCR region of the brain of 0.2 versus 0.04 for the 1.17 mm vs the 1.5mm data, respectively, reflecting the greater uncertainty in determining the fiber orientations in the lower SNR, 1.17mm data; the 0.9mm had about half the SNR of the 1.17mm (35% higher from the longer TE, and less efficient from the longer TR).

For the 1.5mm resolution data with NORDIC processing, approximate gains in fiber orientation dispersion of factor 2 were achievable; for the lower SNR 1.17mm and 0.9mm resolution data, NORDIC yielded larger gains, approximately 5-fold, in fiber dispersion. The *single-resolution multiple repetition data also demonstrated that* gains in fiber orientation dispersion achieved with NORDIC processing correlated with gains in image SNR achieved with averaging multiple acquisitions. Ultimately, the gains realizable in fiber dispersion are probably more limited by the dMRI model fitting, which plateaus at high SNR, as seen with averaging of repetitive scans. This implies that the higher resolution the low SNR is well-suited for locally low rank processing to improve the apparent SNR attained with model-fitting.

The effect of NORDIC is demonstrated on tractography streamlines with two high resolution data in Figure 11. Such streamlines of course do not provide a quantitative demonstration of the effects of NORDIC denoising, but they do provide a *qualitative* and impactful visual demonstration of the improvements achieved with NORDIC denoising. It would not be possible to fully appreciate the extent and the nature of what denoising has done to the dMRI data just by looking at the tractography results, although clearly anatomically well-known tracts are incomplete or absent in the original data and are clearly seen in the denoised data. However, with the detailed and quantitative evaluation presented in Figures 1 through 9, the improvements documented with probabilistic tractography in Figure 10, and tractography streamlines in Figure 11 can *a priori* be expected.

While the presented denoising technique can be applied broadly to other types of image series, the interaction of NORDIC with the model used for assessing the underlying information should be scrutinized. Here, whole brain tractography was used to qualitatively asses the effect of LLR processing, while region-specific analysis was used for quantitative comparison. The conceptual basis of *removal of image content which cannot be distinguished from Gaussian noise* in the LLR model, has broad applications and implications to signal modelling in applications such as ASL, and fMRI and general parameter mapping. In addition to dMRI, ASL and fMRI can also be SNR-starved either due to higher resolution applications and/or the amount of scan time that is available to acquire data – suggesting that such applications could benefit enormously, without cost, from **NORDIC**, and warrants further application specific investigations.

## Conclusion

We propose and validate a noise reduction technique for dMRI using a data processing pipeline that leads to a zero-mean i.i.d. noise component for locally low-rank processing, which enables the use of a parameter-free threshold selection based on random matrix theory. With the *removal of image content which cannot be distinguished from Gaussian noise*, this method was shown to improve model-fitting and structural connectivity mapping. Using NORDIC, the improvements in extractable information content has an increased impact especially for low SNR data, but it would also benefit routine protocols such as the HCP Lifespan.

## Acknowledgement

NIH P41 EB027061, NIH U01 EB025144, NSF CAREER CCF-1651825

