## Supplemental material for "NOise Reduction with DIstribution Corrected (NORDIC) PCA in dMRI with complex-valued parameter-free locally low-rank processing"

Acknowledgement: NIH P41 EB027061, NIH U01 EB025144, NSF CAREER CCF-1651825

For NORDIC processing of dMRI data a two-step (x+t) phase-stabilization is used. The motivation and impact each step is illustrated in supplementary figure S1.

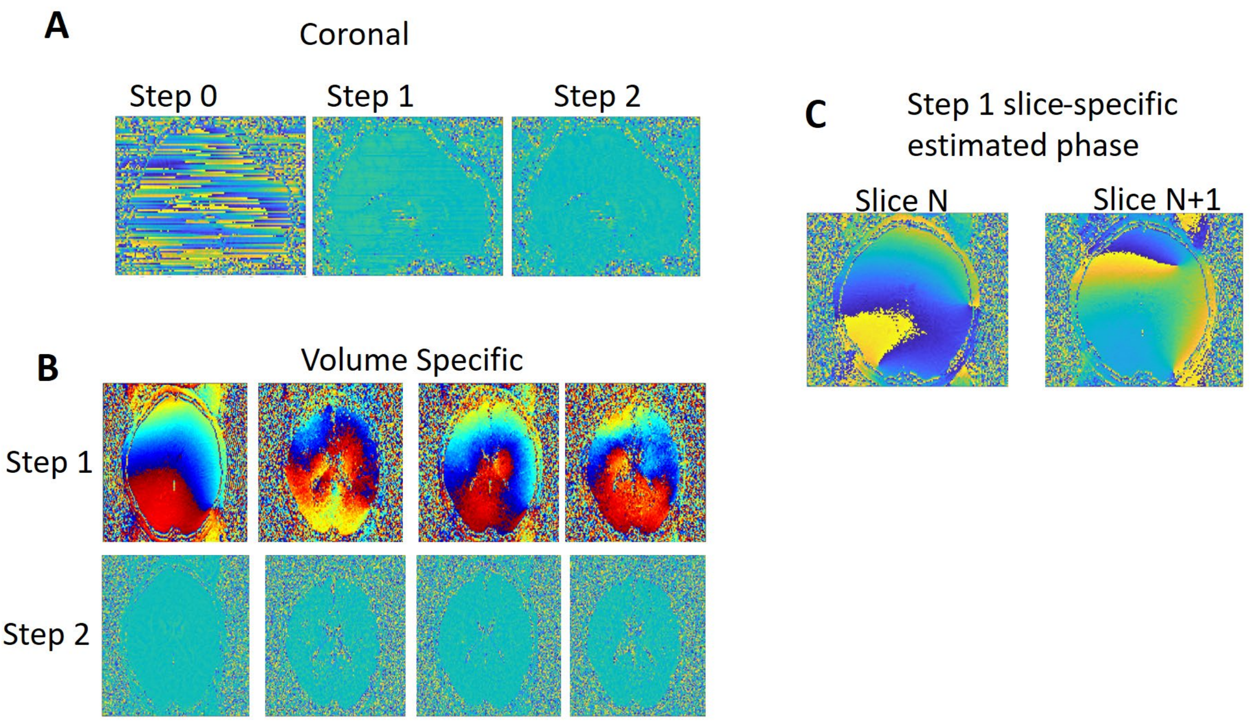


**Figure S1**: The change in phase with the (x+t) phase-correction. Panel A: a coronal cut through the 3D volume is shown, the raw data (step 0, Panel A) exhibit a slice-dependent absolute phase from the estimated sensitivity profiles. This is significantly reduced in Step1 (Panel A), and further suppressed in Step 2 (Panel A). Panel B: top row, the absolute phase in step 1, for 4 different q-vectors is show for an axial slice, and the bottom row show the remaining phase after the correction with step 2. Panel C: the estimated average phase in step 1 is shown for two adjacent slices, reflecting the change necessary.

A real-valued simulation was performed to evaluate the proposed **NORDIC** method using a high SNR reference volume with spatial matrix size 140x140x92, and with 99 volumes (*q*-space samples). Supplemental figure S2 shows the reconstructions and the residuals from the reconstruction.

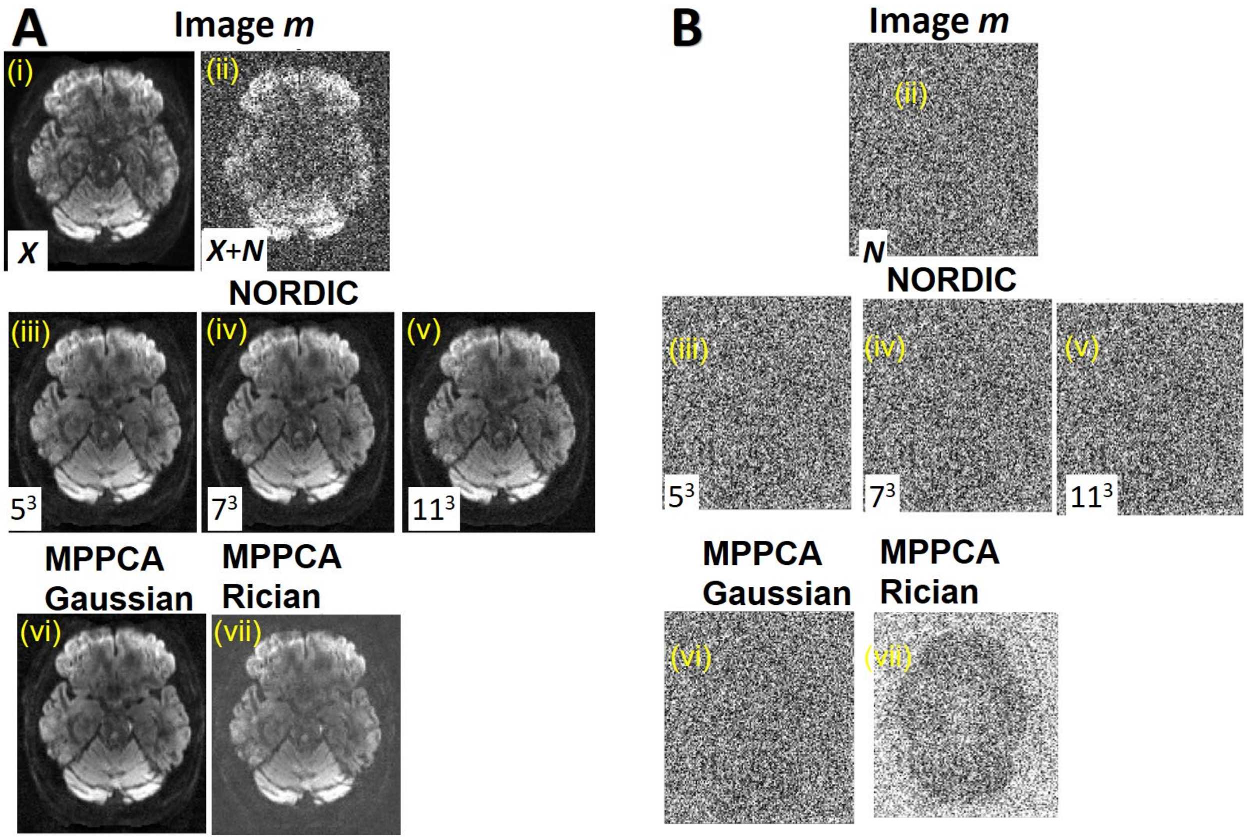


**Figure S2**. Real valued simulation of the quality of the “optimal“ signal recovery with **NORDIC**. **Panel A** shows for a single slice the quality of the images reconstructed with denoising using NORDIC and MPPCA subsequent to SNR degradation with the addition of noise, and **Panel B** shows the difference between the noisy image and the recovered image. The reference images are shown in Fig. S2A.i, before addition of noise and in Fig.S2A.ii, after addition of noise. The NORDIC methods are compared for patch-sizes of $5^3$, $7^3$, and $11^3$ (middle row), and the MPPCA method (bottom row) are compared using the Gaussian noise and Rician noise.

**Figure S3** illustrates an axial slice of the FA map obtained after dMRI processing (right three columns), and the image of the corresponding slice from the volume with b=3000 s/mm$^2$ weighting (left three columns), for the three different resolutions and a single subject. The FA maps and the corresponding slice image with b=3000 s/mm$^2$ are presented for the standard-, MPPPCA and the **NORDIC**-processed data, for the different resolutions. The signal scaling for the bottom row is adjusted for each resolution, since SNR varies with resolution.

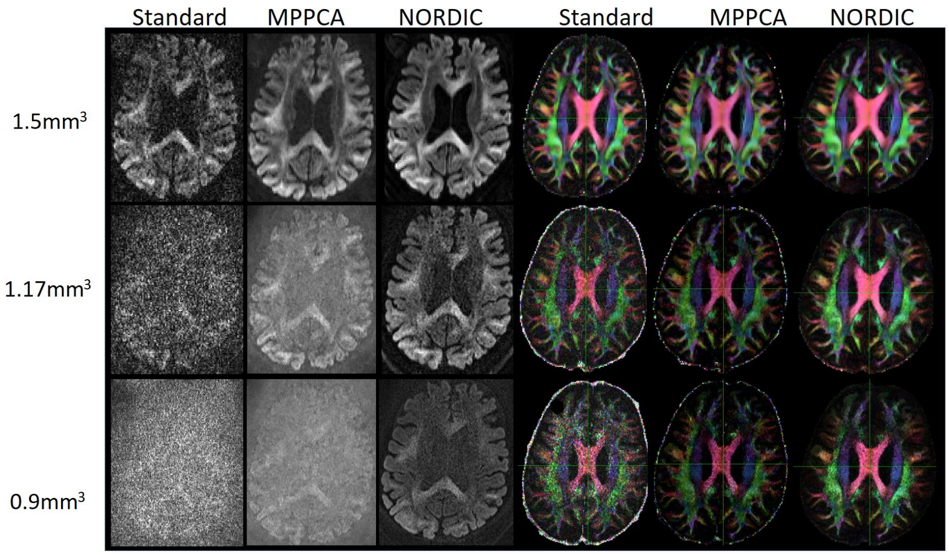


**Figure S3** The effect of NORDIC and MPPCA are shown on a single slice from a diffusion weighted volume (b=3000 s/mm$^2$) across different resolutions (left three columns) and on FA maps (right three columns) for the same slice extracted for the different reconstructions. The FA maps are obtained after EDDY processing, and the reconstructed images are before EDDY processing.

For the 3 subjects with repeated acquisitions, the effect of **NORDIC** and MPPCA after EDDY correction is shown in **Figure S4** for an axial slice in each subject. In this case, each single “repetition” refers to the pair of separate acquisitions with reversed phase encoding which is used for EPI corrections. TOPUP/EDDY, combines data with opposite phase-encoding directions, improving the SNR by approximately $\sqrt{2}$ compared to a true single acquisition.

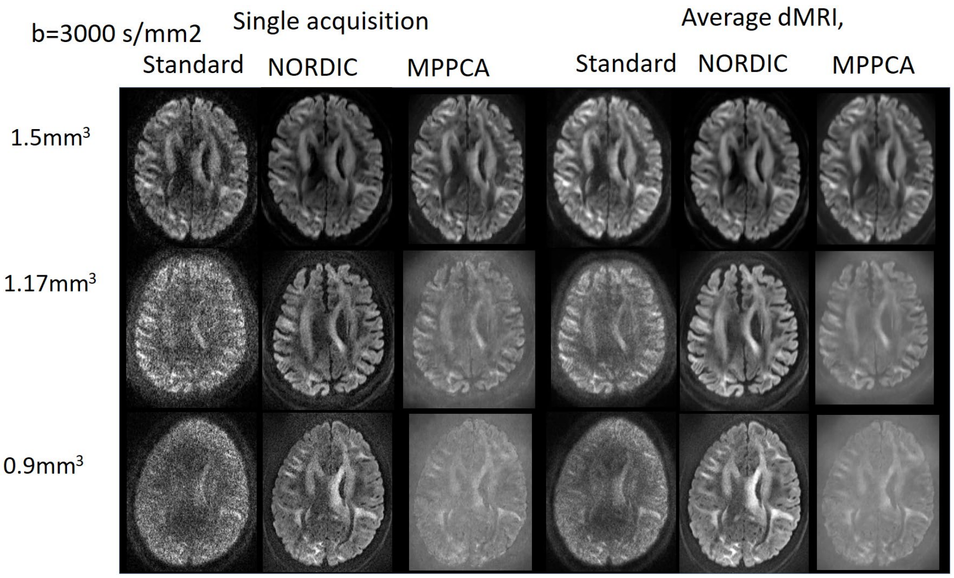


**Figure S4**. Comparison of NORDIC and MPPCA processing with averaging of repetitive acquisitions to increase SNR. The left three columns are for a single acquisition across 3 different resolutions and the right three columns are after averaging 6, 5 and 3 of the repetitive acquisitions respectively. In each case, EDDY processing was applied.

From Figure S5, Panel A, NORDIC processing shows improved connection strength compared to MPPCA processing, which is evident in the corpus callosum area.

At 1.5mm, the difference between MPPCA and NORDIC processed data is less compared to that at higher resolutions, mainly because the number of second fibers estimated at 1.5mm is somewhat similar between these methods, but the improvement in connection strength (the width of the connections shown in yellow) is visible which is due to the lower orientation dispersion and higher number of third fibers resolved in the NORDIC processing.

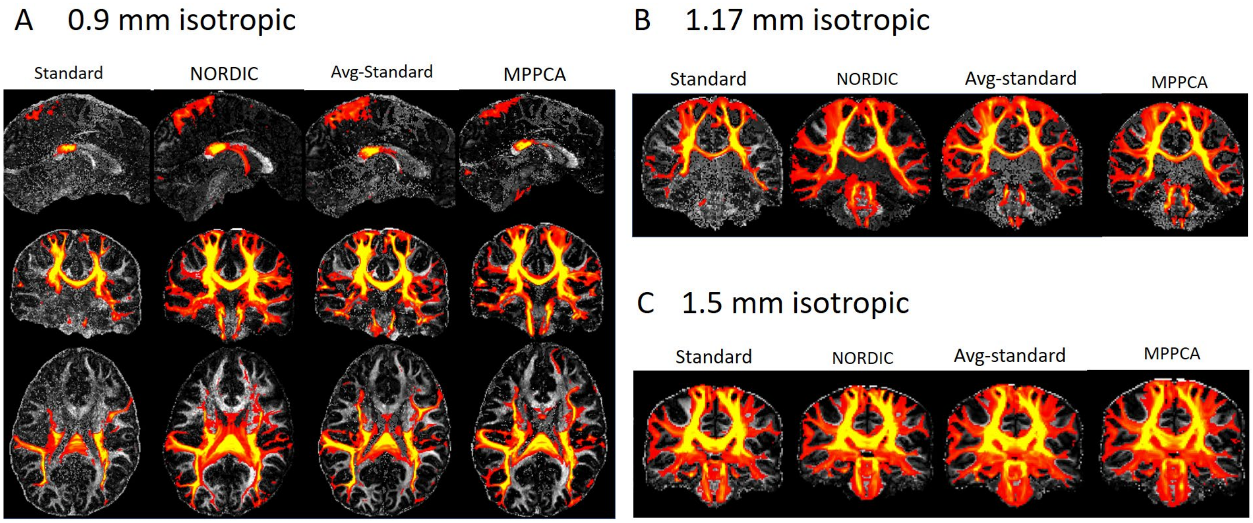


**Figure S5**. Comparison of connectivity distributions from the probabilistic tractography results for 0.9mm (A), 1.17mm (B), and 1.5mm (C) data, representing connectivity of the entire subject-specific posterior corona radiata [PCR]. For each, the figures illustrate the standard reconstruction on a single dMRI data set of the given resolution (labeled as “standard”), standard reconstruction performed on dMRI data obtained with averaging of multiple runs to increase SNR (labeled as Avg-standard), and NORDIC and MPPCA reconstructions of the data without any averaging.

In Supplemental Figure S6 a comparison of denoising with the VST, NORDIC and MPPCA approaches is shown. As the SNR decreases the impact of the non-zero iid thermal noise becomes more apparent for the MPCPA processed data as revealed in a loss of q-space contrast and residual high-spatial frequency modulations in Panel B. The VST algorithm (Ma et al., 2020) confronts the difficulty of working with magnitude data using a two-step approach to resolve the Rician and the spatially varying noise and then uses MPPCA for denoising. Although this approach provides improvements over simply working with magnitude data with Rician noise distribution, it also introduces spatial smoothing which the NORDIC approach does not.

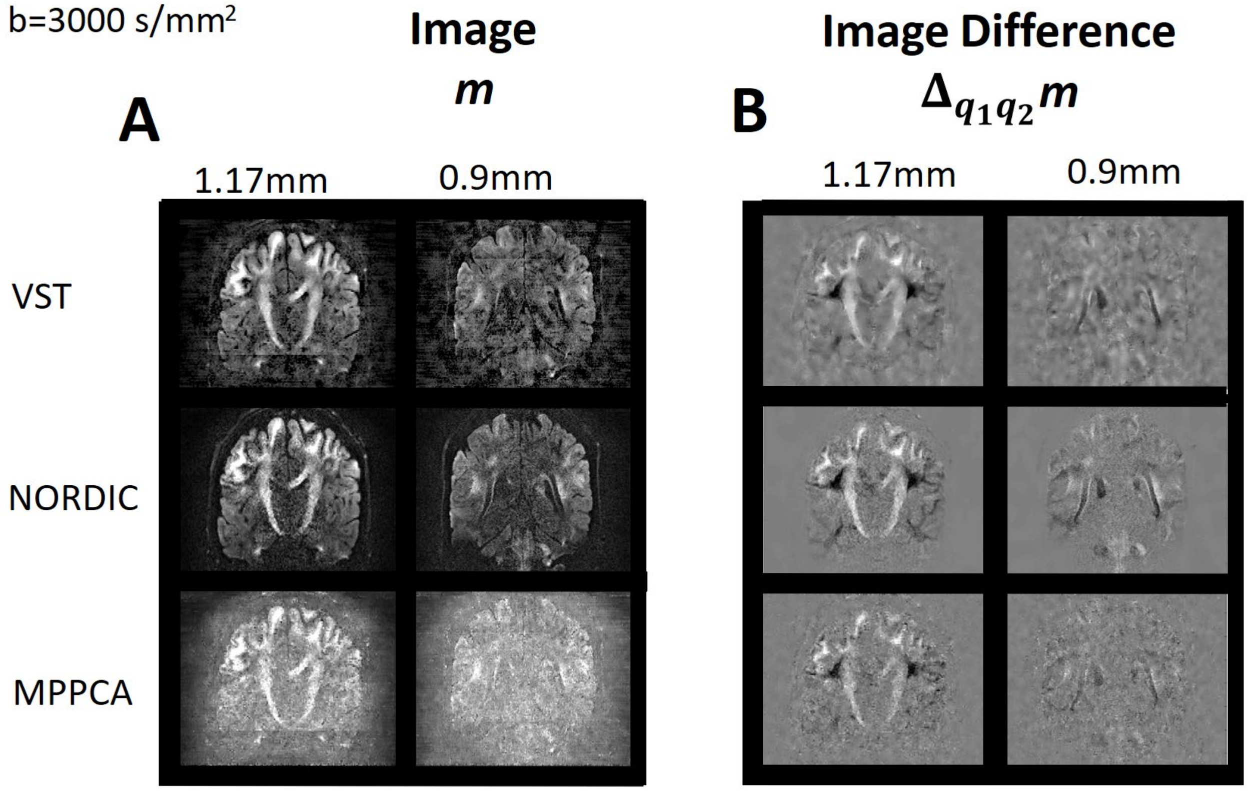


**Figure S6**. Comparison of the denoised images using VST, NORDIC and MPPCA reconstruction. In Panel A, for b=3000 s/mm$^2$, the images are shown for 1.17mm and 0.9mm isotropic resolution. For both VST and MPPCA in Panel A, different residual imaging artifacts are observed, while these are not present in NORDIC. In Panel B, the image difference between two q-space samples with the same b-value are correspondingly shown for VST, NORDIC and MPPCA. For the image difference in Panel B, VST exhibit spatially smoother patches whereas with MPPCA the image difference has more noise relative to both VST and NORDIC. Both VST and MPPCA has lost contrast and features that are preserved with NORDIC. VST in general outperforms MPPCA but not NORDIC.
